# Physical continuity at biomaterial–ECM interfaces regulate fibroblast activation via NF-κB

**DOI:** 10.64898/2026.03.31.715527

**Authors:** Alejandra Suarez-Arnedo, Michaela Harris, Calista Robinson, Lindsay Riley, Amy Kim, Lucy Zhang, Brenton D. Hoffman, Tatiana Segura

**Author notes:** To whom correspondence should be addressed to Tatiana Segura, Brenton Hoffman.

## Abstract

Fibrotic responses at biomaterial–tissue interfaces limit implant integration and regenerative healing, yet how the interaction between biomaterials and the extracellular matrix (ECM) regulates fibroblast activation remains poorly understood. Granular hydrogels including microporous annealed particle scaffolds (MAP) reduce fibrosis, while chemically and mechanically matched hydrogels do not, suggesting a dominant role for scaffold architecture. In this model, MAP scaffolds allow collagen infiltration and form physically continuous composites, whereas hydrogels exclude collagen and generate interfacial slip planes. To isolate how biomaterial architecture influences extracellular matrix (ECM) integration and fibroblast activation, we developed a reductionist in vitro model that integrates collagen type I with either microporous annealed particle (MAP) scaffolds or chemically and mechanically matched bulk hydrogels. This physical integration stabilizes collagen architecture, limits fibroblast-mediated matrix compaction, suppresses contractility, and attenuates myofibroblast transition. Fibroblasts in mechanically integrated environments exhibit reduced expression and nuclear localization of NF-κB and are enriched for quiescent phenotypes. Together, these findings identify biomaterial–ECM physical continuity as a design principle for limiting fibrotic signaling.

## Introduction

Understanding how biomaterials interact with the extracellular matrix (ECM) and resident cells is critical for designing implants that support tissue regeneration while minimizing fibrosis. Fibrotic encapsulation, driven largely by persistent fibroblast activation, compromises implant integration, disrupts tissue mechanics, and limits functional repair [1]. While many biomaterials have been developed to mitigate foreign body responses and promote regenerative microenvironments [2], the physical mechanisms by which biomaterial architectures interact with surrounding ECM to regulate fibroblast behavior remain poorly understood.

Granular hydrogels, particularly microporous annealed particle (MAP) scaffolds [3], have emerged as promising regenerative platforms across applications including wound healing [4, 5], stroke repair [6], and soft-tissue regeneration [7]. Multiple in vivo research demonstrates that MAP scaffolds promote regenerative responses that can prevent fibrotic capsule formation [8], modulate immune responses [9], and lead to tissue repair outcomes more closely resembling native, unwounded tissue [10, 11]. These effects have been associated with multiple design parameters, including pore size [12], packing [13, 14], particle architecture [15, 16], and biochemical functionalization [17, 18]. Collectively, these features influence the behavior and phenotype of infiltrating or encapsulated cells [3, 19], suggesting that MAP scaffold architecture plays an active role in directing tissue regeneration.

Despite these advances, the mechanisms by which the physical architecture of MAP scaffolds contributes to these outcomes remain incompletely understood. While prior studies have largely focused on tunable parameters such as pore size or biochemical cues, less attention has been given to how intrinsic physical features, such as interconnected porosity and scaffold–ECM integration, regulate cell behavior. Notably, MAP scaffolds often outperform conventional hydrogels even when matched in chemical composition and bulk mechanical properties [4], indicating that architecture itself provides distinct physical cues. However, how these architectural features influence the physical and mechanical interactions at the biomaterial–ECM interface, and how these interactions ultimately drive regenerative versus fibrotic responses, remains poorly characterized.

Fibroblasts, which orchestrate ECM deposition, compaction, and remodeling, are highly sensitive to physical and mechanical cues transmitted through the surrounding matrix. Their transition into myofibroblasts is essential for wound repair but can also drive pathological fibrosis when dysregulated [20, 21]. Although biochemical signals such as TGF-β1 are well-known drivers of myofibroblast transition [22, 23], mechanical cues play an equally central role in determining fibroblast phenotype [24–27]. Yet, how these cues differ between granular microporous scaffolds and hydrogels, and how such differences influence fibroblast transition, remains unclear.

Here, we hypothesized that the granular architecture of MAP scaffolds modulates the surrounding ECM, altering collagen assembly and physical continuity. Physical continuity is defined here as a seamless integration between the biomaterial and surrounding ECM, and these changes act to control fibroblast transition. To test this hypothesis, we developed a reductionist, in vitro model integrating collagen type I matrices with either MAP scaffolds or chemistry-matched hydrogels. We observed fundamental differences in how these materials interact with the surrounding matrices and affect cell behavior. Specifically, we find that granular MAP architectures enable physical scaffold–matrix integration, which functions as a potent inhibitor of fibroblast activation. We suggest that ensuring mechanical connectivity between biomaterial implants and surrounding ECMs is a key design principle for preventing fibrosis and supporting functional tissue regeneration.

## Results

### Collagen fibers assemble within the void spaces of MAP scaffold to promote physical integration and minimize slip plane formation observed in hydrogels

To examine how ECM assembles in the presence of biomaterials, we developed an in vitro assay that enables visualization of collagen fibrillogenesis surrounding MAP scaffolds or hydrogels that are chemically and mechanically similar (Figure 1A, Figure 2A). Both materials were fabricated from the same 8-arm polyethylene glycol–vinyl sulfone (PEG-VS), RGD adhesion motifs, and crosslinked with an MMP-labile. This formulation was selected because the same MAP formulation has previously demonstrated regenerative outcomes in vivo [9]. In this design, the MMP-labile crosslinker provides enzymatic degradability [28], while RGD gives integrin-binding sites that support cell adhesion [29]. Rheological characterization confirmed that MAP scaffolds and bulk hydrogels display comparable bulk storage and loss moduli (Figure 1B, C, and G), indicating similar macroscopic mechanical properties. Importantly, prior work has also demonstrated that microgels fabricated from identical chemistry possess equivalent local Young’s modulus to bulk gels of the same formulation, confirming that the particle building blocks of MAP scaffolds share the comparable stiffness as their bulk counterparts [13]. Together, these results indicate that any differences in collagen assembly arise from the structural organization of the materials rather than from variations in chemistry or mechanics. For this acellular model, collagen polymerization was carried out at 4□°C to slow fibrillogenesis and promote formation of well⍰defined fibers [30–32]. Under these conditions, the fibrils interweave into a dermis⍰like meshwork while maintaining bulk stiffness comparable to gels polymerized at higher temperatures [32], yielding a physiologically relevant ECM structure that is also compatible with high⍰resolution imaging and quantitative analysis. Similarly, centrifugation was applied prior to fibrillogenesis to facilitate uniform infiltration of the collagen solution into the scaffold’s interstitial spaces, ensuring homogeneous filling of the pores (Supplementary Figure□1) and preventing quantification artifacts associated with voids or uneven fiber assembly. This procedure produced collagen networks that closely resemble histological sections in vivo, where there are no large fluid⍰filled gaps between granular biomaterials and the surrounding ECM [9].

**Figure 1.**
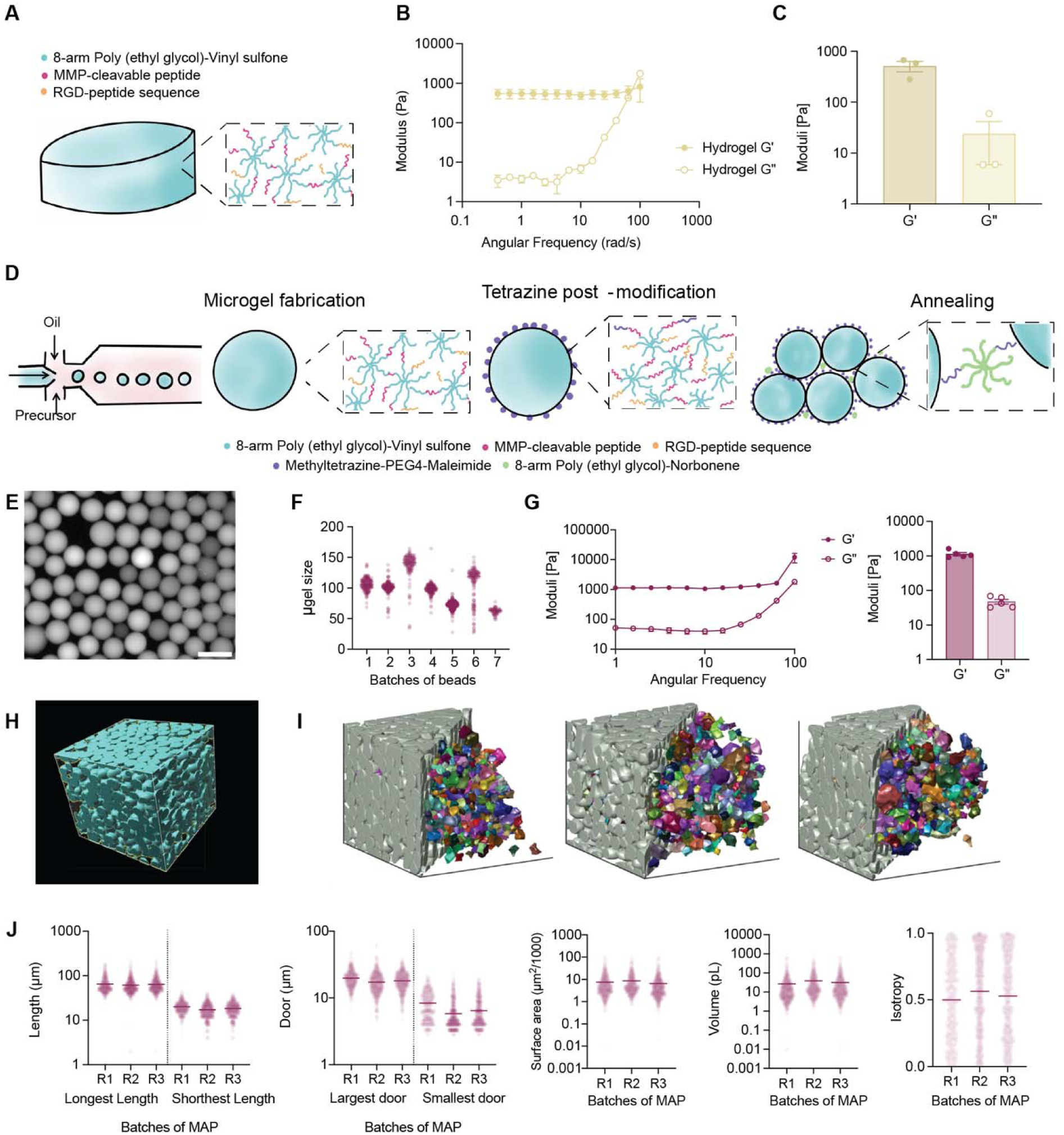
Using MAP Scaffolds and Hydrogels with Matched Chemistry and Mechanical Properties. (A) Schematic representation of PEG hydrogel chemistry using Michael Addition. (B) Frequency sweep of PEG hydrogels at 1% strain; data points represent the mean ± SD from three independent hydrogel batches (N = 3). (C) Storage and loss moduli (G’, G’’) of PEG hydrogels, with each bar showing the mean ± SD from three independent batches (N = 3). (D) Schematic representation of microgel composition and post-conjugation chemistry for norbornene-tetrazine annealing. (E) Representative image of microgels. (F) quantification of multiple batches of microgels used. Scale bar = 200 µm. (G) MAP scaffold Frequency sweep, strain % 1 and Storage and Loss modulus of MAP. N = 3 independent batches of MAP scaffold (H) 3D rendering of MAP scaffold. (I) LOVAMAP images showing the MAP scaffold (left) and its interior pores (right) (J) length, door diameter, surface area, volume and isotropy (i.e., orientation) of the pores using LOVAMAP. All data are presented as averages ± S.E.M. Statistical significance was determined using a Nested ANOVA with Dunnett’s post-hoc multiple comparison test (*p < 0.05; N = 3 independent batches of MAP).

**Figure 2.**
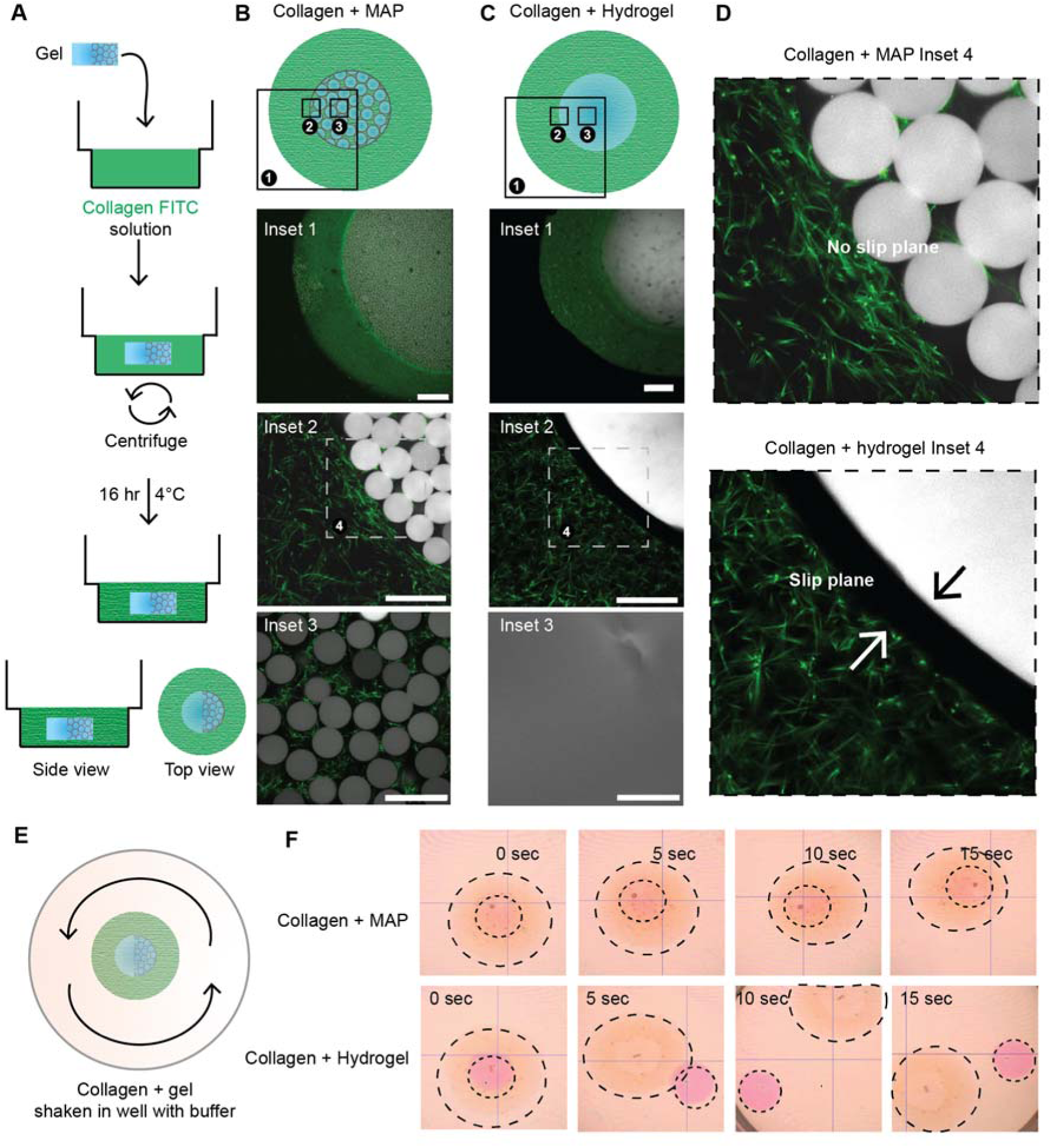
MAP gels supported ECM assembly throughout their structure, while a clear slip plane was observed between the ECM and traditional hydrogels. (A) Schematic of experimental setup for introducing FITC-labeled collagen into MAP scaffolds or hydrogels, followed by centrifugation and incubation. (B) Confocal images of MAP scaffolds show uniform collagen integration between microgel voids (Insets 1–3), with no detectable interface or slip plane formation. (C) In contrast, collagen interfacing with hydrogels results in clear exclusion from the hydrogel compartment, forming a sharp, persistent slip plane at the boundary (Insets 1–3). (D) Magnified views highlight absence of a slip plane within MAP scaffolds (upper), but clear slip plane (arrows) at the hydrogel boundary (lower). (E) Illustration of movement assay evaluating detachment at the interface upon shaking. (F) Sequential images show collagen-MAP constructs retain integration during movement, while collagen-hydrogel constructs display separation and movement of the gel interface over time, consistent with slip plane formation. Scale bars: Scale bars: 500 μm (inset 1); 200 μm (Inset 2)

When collagen was introduced to each biomaterial, we observed strikingly different collagen assembly behaviors. Despite their identical chemical and mechanical properties, collagen fibers readily assembled outside and throughout the interconnected void spaces of MAP scaffolds, forming continuous fibrillar networks that spanned neighboring microgels (Figure 2B–D). In contrast, collagen interfacing with hydrogels resulted in collagen assembly out of the biomaterial but was clearly excluded from the hydrogel compartment. This produced a sharp and consistent slip plane at the boundary. These findings indicate that the microporous architecture of MAP scaffolds governs their ability to integrate with the ECM.

To further understand the physical features that enable this integration, we characterized MAP scaffold architecture using LOVAMAP, an image-analysis tool for identifying pores within granular materials and for measuring features like pore size, orientation, and width of openings (i.e., “doors”) [33] between adjacent pores (Figure 1H–J) [34]. This analysis revealed pores averaging 60 µm along their longest axis and with 5–20 µm door diameters, formed by microgels approximately 100 µm in diameter. These pore dimensions are sufficiently large to accommodate both collagen fibers (diameter ≈ 2 µm, length ≈ 5–13 µm) and cells such as fibroblasts with reported diameters around 18 µm [35], establishing a microporous network capable of hosting both ECM fibers and cells. In contrast, the nanoscale mesh of hydrogels (∼18 nm) physically restricts collagen fiber assembly. Overall, these findings suggest that the larger, connected pores in MAP scaffolds enable continuous collagen and cell networks across the biomaterial interface, whereas the nanoscale pores in bulk hydrogels limit physical connectivity and therefore mechanical integration.

To assess whether this physical continuity translated to mechanical integration, we applied controlled agitation to the collagen–biomaterial composites (Figure 2E). Collagen–MAP composites remained integrated during movement, whereas collagen–hydrogel composites rapidly detached at the interface (Figure 2F). The detachment of the hydrogel core demonstrated the presence of a mechanical slip plane due to the loss of physical continuity between the ECM and the hydrogel. In contrast, the seamless interpenetration of collagen fibers within MAP scaffolds prevented interfacial separation, producing a mechanically integrated composite.

Together, these findings demonstrate that collagen fibers assemble within the void spaces of MAP scaffolds, forming continuous networks that bridge the scaffold and surrounding ECM. The granular MAP design enables mechanical integration that prevents slip plane formation. In contrast, hydrogels, with their homogenous, nanoporous structure, fail to integrate physically with the ECM, leading to separation under stress.

The presence of the granular scaffold alters collagen fiber architecture at the scaffold interface, creating a distinct edge effect at adjacent regions.

Using the same in vitro assay, we next investigated how the presence of these biomaterials influences collagen fiber organization at their interfaces. Collagen matrices were polymerized around either MAP scaffolds or hydrogels, and confocal microscopy combined with IMARIS-based filament tracing was used to quantify fiber morphology and distribution near the biomaterial boundaries (Figure 3A, Supplementary Figure 2). Within the MAP condition, collagen fibers were observed to densely accumulate at the scaffold interface, forming a high-intensity band that extended several micrometers into the surrounding matrix (Figure 3B). Quantitative analysis revealed that fibers within this region displayed increased length, volume, and local density relative to fibers farther from the scaffold surface. This localized increase in collagen fiber metrics produced a distinct and statistically significant edge effect at the MAP–collagen interface, consistent across independent replicates (Figure 3B). Despite this enhanced local density, collagen fibers retained their isotropic orientation, suggesting that MAP scaffolds promote collagen matrix integration without inducing large-scale directional alignment of the collagen fibers. Within the hydrogel condition, collagen matrices polymerized adjacent to the hydrogels showed no such edge effect, with collagen fibers showing a uniform distribution across the collagen matrix (Figure 3C). Together, these results further support the finding that collagen-MAP scaffolds produce a mechanically stable composite and do so through enriching collagen fiber assembly at the collagen-MAP interface, leading to an edge effect. Conversely, hydrogels do not support localized fibrillogenesis at the collagen-hydrogel interface, causing the collagen matrix to form independently of the hydrogel matrix and resulting in a slip plane between the collagen matrix and the hydrogel.

**Figure 3.**
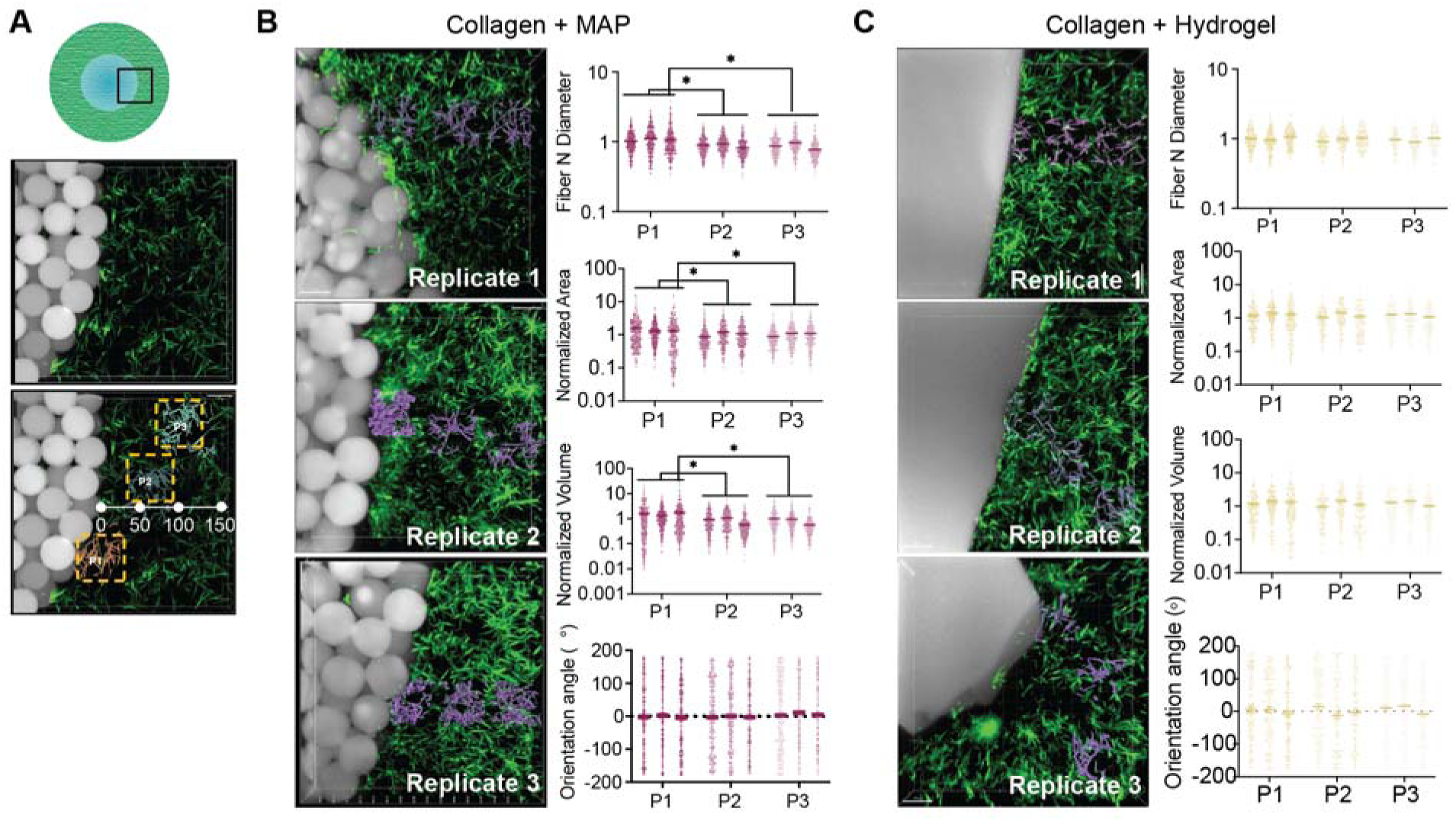
MAP scaffolds in collagen matrices alter neighboring fibril formation. **(A)** (A) Schematic and representative region-of-interest showing analysis zones at the MAP–collagen interface. (B) Confocal images and quantification of collagen fibers adjacent to MAP scaffolds reveal high fiber density, area, and volume with significant batch variability across three replicates, demonstrating robust local matrix assembly. (C) At hydrogel–collagen interfaces, confocal images show lower and more uniform fiber metrics across replicates, indicating an absence of local matrix enrichment. All data are presented as averages ± S.E.M. Statistical significance was determined using a Nested ANOVA with Dunnett’s post-hoc multiple comparison test (*p < 0.05; N = independent batches of collagen gels). Scale bars: 40 µm

### MAP reduces fibroblast-mediated compaction in collagen gels

We next investigated whether the differences in collagen integration in MAP versus hydrogel affect the ability of fibroblasts to compact collagen We used a collagen gel compaction assay, in which fibroblasts are seeded in a collagen gel for 48□h and subsequently released from the well edge to allow collagen contraction [36]. In this study, we specifically employed the stressed-matrix compaction variant of this assay, where the collagen gel is first constrained during culture and then released after several days to quantify fibroblast⍰driven remodeling [37]. Collagen-containing fibroblasts were cast in a 48-well plate, and MAP or hydrogel was placed in the collagen solution before fibrilization (Figure 4A). As a result, each composite gel contained MAP or hydrogel embedded within the collagen–fibroblast matrix. Cells were cultured in the composite gel for 48-hours before collagen compaction was initiated by releasing the gels from the well edges for 5 hours (Figure 4A). Because fibroblasts are known to reorganize collagen and generate tensile forces that compact the matrix [36, 38], this assay enables direct comparison of how scaffold integration affects cell-driven mechanical remodeling. This approach also allowed us to test whether the mechanical integration and edge-effect observed in acellular systems (Figures 2–3) would occur under conditions where the matrix is actively remodeled by fibroblasts.

**Figure 4.**
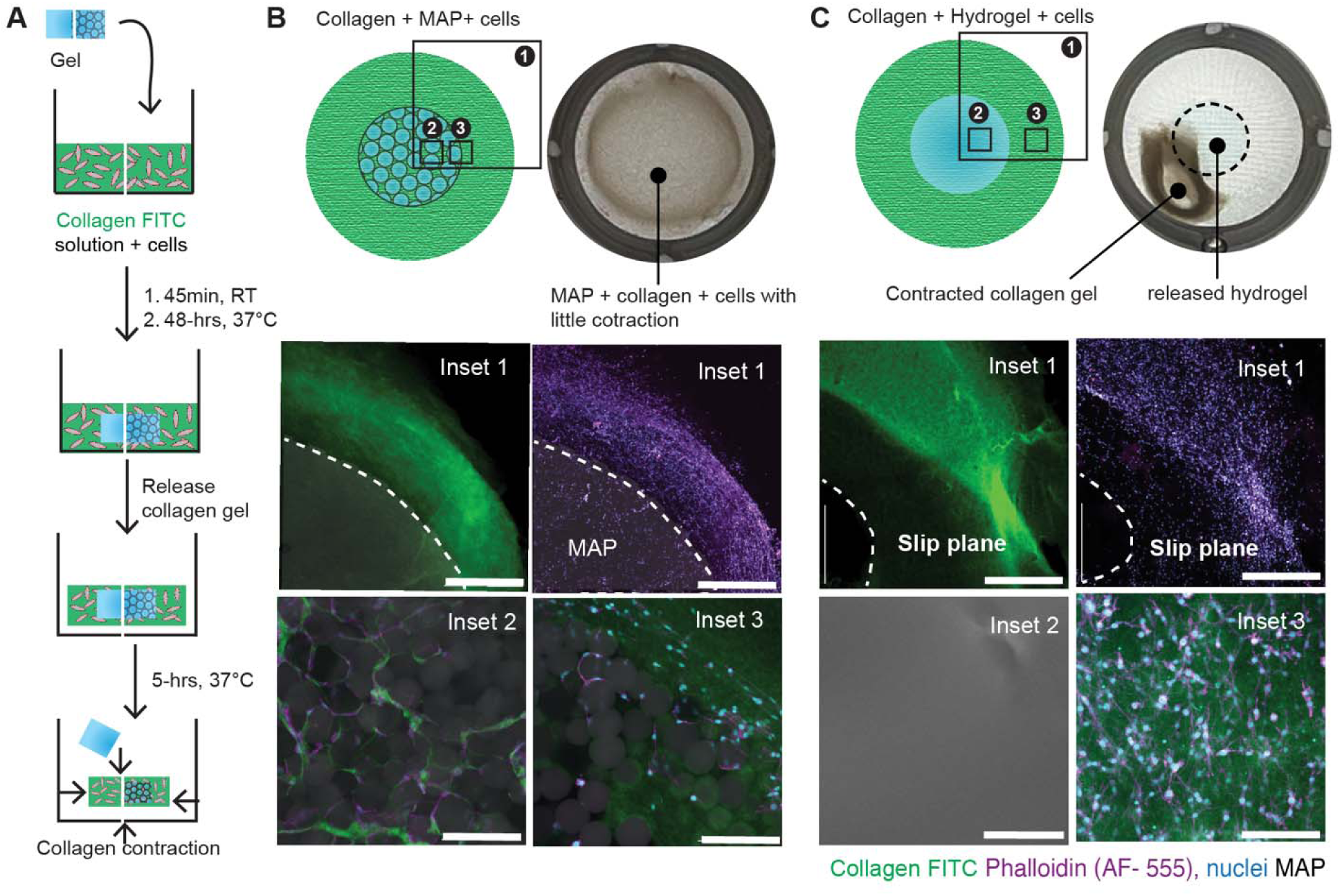
MAP scaffolds integrate seamlessly with collagen matrices in cellular systems, whereas hydrogels form a slip plane that prevents integration. **(A)** Experimental schematic showing encapsulation of cells within FITC-labeled collagen in contact with either MAP scaffolds or hydrogels, followed by incubation, gel release, and compaction. (B) Collagen + MAP scaffolds demonstrate uniform cell and collagen infiltration with little contraction and no slip plane at the interface (Insets 1–3). Confocal images confirm MAP is well integrated with the surrounding collagen and cells. (C) In contrast, collagen + hydrogel constructs exhibit pronounced collagen gel contraction and release of the hydrogel core. Confocal imaging reveals a distinct slip plane at the hydrogel–collagen interface, visualized by sharp boundaries and exclusion of cells and collagen fibers (Insets 1–3). Channel assignments: collagen (green, FITC), actin (magenta, AF-555 phalloidin), MAP (grey, AF-647 maleimide), and nuclei (blue). Scale bars: Scale bars: 500 μm (inset 1); 200 μm (Inset 2 and 3).

We first investigated whether collagen and cells were observed inside either MAP or hydrogel. In collagen + fibroblast + MAP constructs, fibroblasts and collagen fibers were observed in the interconnected microgel network, while in the collagen + fibroblast + hydrogel constructs, neither collagen nor fibroblasts were observed inside the hydrogel. Surprisingly, even after collagen detachment from the well plate, the collagen + fibroblast + MAP condition did not show significant contraction of the collagen matrix (Figure 4B). In contrast, collagen + fibroblasts + hydrogel constructs exhibited pronounced global compaction accompanied by rapid detachment of the hydrogel from the surrounding matrix (Figure 3C). These results suggest that the physical continuity observed in collagen + MAP scaffolds maintains a structurally integrated composite even under cell-generated forces, while the collagen + hydrogel composites remain structurally separated through a slip plane, which allowed significant collagen contraction under cell-generated forces. These findings demonstrate that MAP scaffolds and hydrogels create fundamentally different mechanical environments for fibroblasts.

We next examined how the presence of cells influences the organization of collagen in the composite hydrogels. Regions adjacent to MAP scaffolds retained the dense, spatially localized collagen enrichment characteristic of the acellular edge effect, with intensity returning to baseline within ∼10 μm of the microgel surface (Figure 5B–C). This indicates that MAP scaffolds preserve their interfacial microarchitecture even during fibroblast remodeling. In contrast, collagen + hydrogel constructs developed broad (>100 μm) regions of elevated collagen density (Figure 5D), reflecting the compaction and slip plane seen at the macroscopic scale. Together, these observations show that MAP scaffolds stabilize the local collagen architecture, whereas hydrogels promote heterogenous matrix organization.

**Figure 5.**
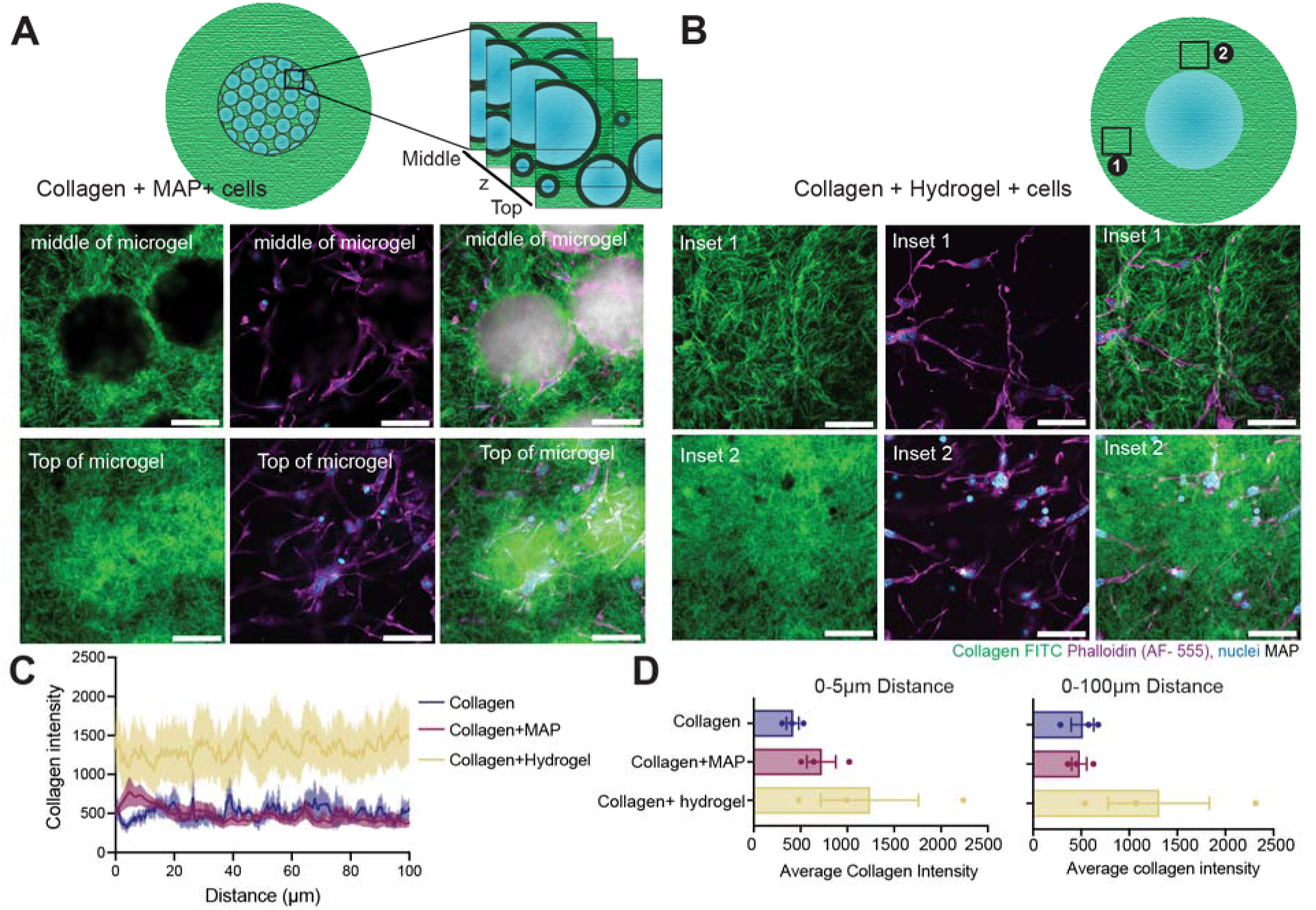
Collagen fiber density remains elevated near MAP scaffold surfaces even in cellular models. (A) Schematic of imaging regions and representative confocal slices from the middle and top of MAP microgels, showing high local collagen (green, FITC) density. (B) In hydrogels, confocal images from different regions reveal collagen increases fiber density, (C) Quantitative profiles of collagen intensity as a function of distance from the interface compare collagen-only, MAP-collagen, and hydrogel-collagen constructs. (D) Collagen density within 0–5 μm and 0–100 μm . All data is represented by average ± S.E.M. N = Number of collagen gels, each containing cells from independent cultures. Scale bars: 20 µm.

We next moved to further characterize fibroblast-mediated collagen compaction. Fibroblast-driven gel compaction is a classic metric of fibroblast transition into myofibroblast and matrix–cell mechanical coupling [39] and is highly sensitive to biochemical cues, ECM rearrangement, and cell contractility [36, 40] and biomaterial integration. As already mentioned, we adapted a collagen gel compaction assay to evaluate whether the presence of a MAP scaffold or a hydrogel modulates the contractile behavior of fibroblasts embedded within collagen (Figure 6A). TGF-β1 was used as a stimulatory signal for fibroblast transition to myofibroblasts.

**Figure 6.**
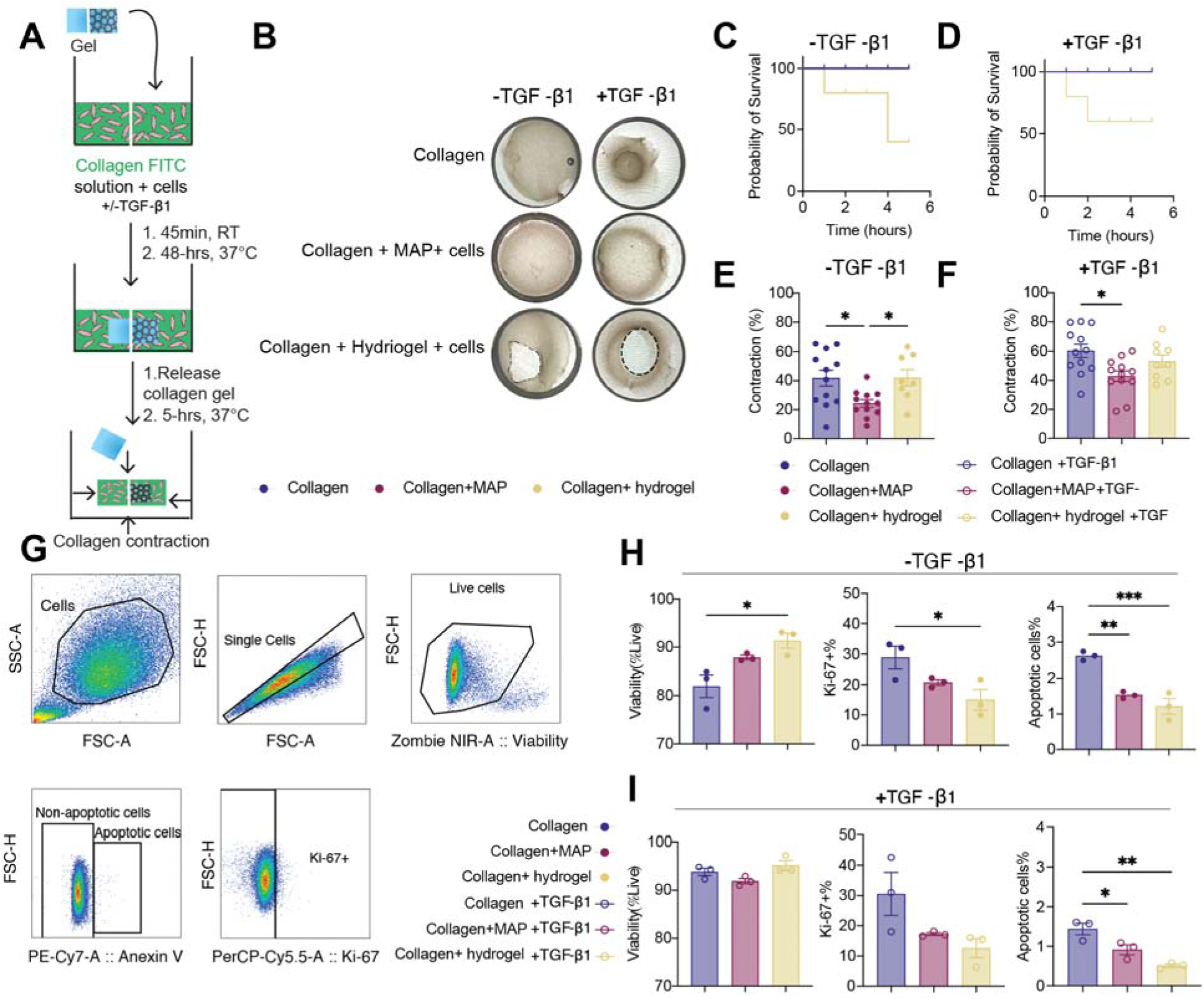
MAP scaffold presence reduces collagen compaction. (A) Schematic of the collagen gel compaction assay workflow. (B) Representative compaction images for collagen-only, collagen + MAP, and collagen + hydrogel constructs, with and without TGF-β1 treatment. (C, D) Survival curves indicate gel stability under each condition. (E, F) Quantification of contraction reveals that MAP scaffolds minimize compaction compared to hydrogel and collagen-only controls, regardless of TGF-β1 addition. (G) Flow cytometry strategy for identifying live, apoptotic, and proliferating (Ki-67+) cells. (H, I) Comparative analysis of fibroblast viability, apoptosis, and proliferation (Ki-67+) across collagen matrices type, of fibroblasts activated with TGF-β1. Data are presented as mean ± S.E.M. Filled circles represent measurements from samples treated without TGF-β1, while open circles correspond to samples treated with TGF-β1. Statistical analysis performed using one-way ANOVA with Tukey’s post-hoc test (*p<0.05, **p<0.01, ***p<0.0001). N = Number of collagen gels, each with independent cell culture.

Visual inspection revealed that collagen + MAP matrices exhibited uniform compaction without loss of MAP–collagen continuity, consistent with the physical integration observed in acellular systems during shaking. By contrast, collagen + hydrogel constructs demonstrated pronounced detachment of the hydrogel core during contraction, consistent with core detachment during agitation of the acellular system (Figure 6B, Supplementary Figure 3). To quantify these hydrogel⍰core detachment events, we used a survival curve in which gels exhibiting core detachment was scored as non-surviving. This analysis revealed a higher probability of failure for hydrogels compared to non-failures in MAP constructs (Figure 6C, D). These spontaneous core⍰detachment events reflect failure at the slip plane and further support the finding that the hydrogel is not mechanically coupled to the surrounding collagen. As expected, TGF-β1 increased the compaction rate observed in collagen-only gels; however, for the composite gels, the effect of TGF-β1 addition was less pronounced (Figure 6B).

Quantitatively, collagen-only matrices compacted steadily, reaching an average of 40% contraction at 5 hours (Figure 6E), and as expected, contraction was further increased by TGF-β1 stimulation, reaching an average of 60% contraction at 5 hours (Supplementary Figure 3A, B, C). Collagen + MAP composites markedly reduced collagen gel contraction relative to both collagen-only and collagen + hydrogel conditions (Figure 6E–F). Specifically, MAP-containing gels compacted an average of 20% without TGF-β1 and 40% with TGF-β1 (Supplementary Figure 3A-C). The collagen + hydrogel condition was the only condition that did not have a statistically significant increase in contraction when TGF-β1 was added but achieved significant contraction in the absence of TGF-β1 (Figure 6E, Supplementary Figure 3A, B, C). Together, these findings show that MAP scaffolds dampen fibroblast⍰driven matrix remodeling, as reflected by the reduced gel contraction compared with collagen⍰only controls.

We next assessed cell viability, apoptosis, and proliferation following collagen compaction using flow cytometry. After 5 hours of contraction, cells were released from the composite material using Liberase, stained, and analyzed with flow cytometry. Consistent with previous reports that TGF-β enhances fibroblast viability by reducing apoptosis [22, 41, 42], TGF-β1 treatment increased viability and lowered the apoptotic fraction across all conditions (Supplementary Figure 3D). In addition, the presence of a biomaterial, whether MAP or hydrogel, reduced Ki-67⁺ proliferation relative to collagen-only controls (Fig. 6H–I), indicating that biomaterials locally reduce fibroblast proliferation in 3D collagen environments, in line with prior findings that ECM interfaces modulate fibroblast growth through mechanical and biochemical cues [27, 43]. Importantly, differences in apoptosis between MAP and hydrogel conditions were modest, demonstrating that the divergent functional outcomes (reduced compaction in MAP composites versus heightened contraction in hydrogel constructs) cannot be attributed to differences in cell number or survival. Instead, these results support a model in which the physical continuity provided by MAP scaffolds attenuates myofibroblast contractility, whereas slip-plane formation in hydrogel constructs generates a e microenvironment that increases collagen compaction (Fig. 6B–F).

### Collagen + MAP composite limits fibroblast activation, decreasing collagen and NFκB expression associated with the activated phenotype

We next investigated whether the structural and mechanical differences translate into changes in fibroblast phenotypic state. Fibroblast transition into a myofibroblast phenotype is characterized by increased contractility, ECM production, and pro-fibrotic phenotype, and is regulated by both biochemical cues (e.g., TGF-β) and mechanical inputs from the surrounding matrix [20, 40, 44]. Because this transition is accompanied by well-defined molecular changes, we selected markers that together capture cytoskeletal activation (αSMA) [45, 46], matrix synthesis (Collagen I) [46, 47], mechanotransduction (YAP) [48, 49], and inflammatory/pro-fibrotic phenotype (NFκB) [50–52]. To address this, we performed immunofluorescence imaging of these key markers associated with fibroblast transition and pro-fibrotic phenotype, using identical collagen matrices containing either no biomaterial, a MAP scaffold, or a hydrogel (Figure 7A–B).

**Figure 7.**
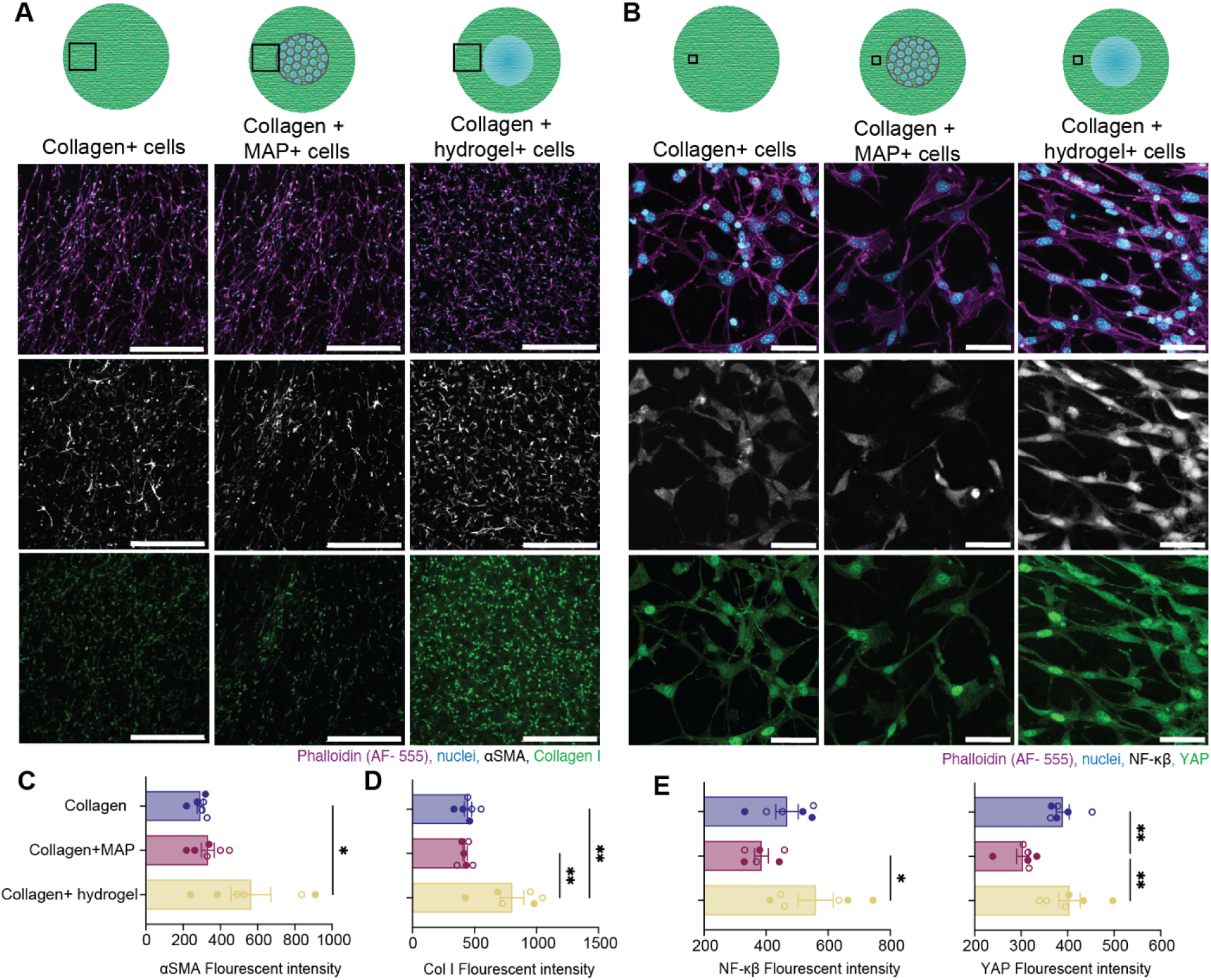
Cellular activation and matrix synthesis are modulated by biomaterial presence. (A) 20X Representative confocal fluorescence images of cell-laden collagen gels, collagen + MAP, and collagen + hydrogel matrices. Panels show composite and single-channel images for actin filaments (phalloidin, magenta), nuclei (blue), αSMA (white), collagen I (green) as indicated. Schematics above each panel illustrate experimental setup and sampling regions. Marker expression and cellular morphology are compared for each material interface. (B) 60 X Representative confocal fluorescence images of cell-laden collagen gels, collagen + MAP, and collagen + hydrogel matrices. Panels show composite and single-channel images for actin filaments (phalloidin, magenta), nuclei (blue), NF-κB (white), and YAP (green), as indicated. Schematics above each panel illustrate experimental setup and sampling regions. Marker expression and cellular morphology are compared for each material interface. (C-F) Quantification of αSMA, collagen I, NF-κB, and YAP fluorescence intensities per area across all conditions demonstrates differences in cellular activation and matrix remodeling. Data are presented as mean ± S.E.M. Filled circles represent measurements from samples without TGF-β1, open circles correspond to samples with TGF-β1. Statistical analysis was performed using one-way ANOVA with Tukey’s post-hoc test (*p<0.05, **p<0.01, ***p<0.001, ****p<0.0001). N = Number of collagen gels, each with independent cell culture.

Notably, TGF-β1 stimulation did not significantly change αSMA, Collagen I, YAP, or NFκB expression across any matrix condition (Supplementary Figure 4). This finding is consistent with known effects of mechanically prestressed collagen matrices, in which matrix tension alone can sustain fibroblast transition independent of added TGF-β1 [37, 53]. Because TGF-β1 did not alter the relative trends among biomaterial conditions, data from treated and untreated samples were pooled for subsequent analysis. Additionally, across all measurements of marker expression, responses in collagen + MAP matrices did not vary between regions adjacent to the microgels and regions farther away (Supplementary Figure 5), indicating that any transition profile is long-range across the gel rather than restricted to the scaffold interface. This is consistent with the absence of localized, heterogeneous compaction zones in MAP composites and suggests that maintaining mechanical integration prevents the development of localized stress fields that promote fibroblast activation [43].

Across all markers, collagen + MAP matrices consistently exhibited lower levels of fibroblast transition relative to collagen-only and collagen + hydrogel controls (Figure 7C–F). Fibroblasts on collagen + MAP matrices displayed reduced αSMA expression, indicating attenuation of the contractile myofibroblast phenotype. Similarly, Collagen I deposition, another hallmark of fibroblast transition [54], was significantly reduced in MAP-containing gels. In contrast, fibroblasts in collagen + hydrogel constructs exhibited elevated Collagen I and αSMA expression, consistent with a more pro-fibrotic phenotype. These patterns were observed despite similar or lower levels of proliferation in the hydrogel condition (Fig. 6H–I), indicating that increased collagen expression in the hydrogel system arises from altered cell state rather than increased cell number.

We next examined two signaling proteins associated with mechanotransduction (YAP) and inflammation-driven fibrosis (NFκB) [49, 51, 52]. Because both pathways are regulated not only by overall expression but also by stimulus-dependent nuclear translocation, we quantified total protein levels as well as nuclear-to-cytoplasmic (N/C) ratios to capture changes in phenotype state [52, 55–57]. Fibroblasts in collagen + MAP matrices displayed minimal NFκB expression and low YAP levels, suggesting reduced inflammatory and mechanical pathway activation. In contrast, collagen + hydrogel constructs showed markedly higher NFκB and YAP expression (Figure 7B, 7E–F). The hydrogel condition also showed increased nuclear localization of NFκB, and similar nuclear translocation of YAP compared with MAP and collagen-only conditions (Supplementary Figure 7). Because YAP nuclear translocation was elevated across all matrix types, including collagen-only, this pattern is likely driven by the substantial tensile forces generated during the collagen compaction rather than by biomaterial-specific effects, making it difficult to disentangle the contributions of collagen-generated forces from any influence of the biomaterial itself. This interpretation is further supported by prior reports demonstrating that increased matrix tension and mechanical prestress are sufficient to drive YAP nuclear translocation, even in the absence of additional biochemical cues [43, 55]. Given this, and because NFκB nuclear enrichment is a well-established feature of inflammatory fibroblasts [56], we focus our subsequent analyses on the expression and nuclear localization patterns of NFκB. These observations support the broader trend that collagen + hydrogel matrices promote an inflammatory/myofibroblast-like phenotype, whereas MAP scaffolds attenuate fibroblast transition.

To determine whether NFκB nuclear translocation correlated with the degree of compaction (Figure 6), we quantified NFκB nuclear/cytoplasmic ratios across gels. After compaction, collagen + hydrogel matrices exhibit the highest NFκB nuclear localization, while collagen + MAP matrices maintain low levels comparable to collagen-only controls (Supplementary Figure 7). These results indicate that the hydrogel⍰core detachment and compaction in hydrogel constructs selectively trigger NFκB translocation, whereas the mechanically integrated MAP matrices suppress compaction-induced inflammatory activation.

To assess whether NFκB or YAP activation could be explained by changes in cell shape, spreading, or cytoskeletal structure, we correlated NFκB or YAP expression and nuclear translocation with morphological metrics such as cell area, aspect ratio, and actin organization. Although previous studies have shown that NFκB nuclear translocation can be influenced by cell shape [58], cytoskeletal tension, and mechanotransduction pathways [59], and that YAP localization is similarly regulated by mechanical cues and cytoskeletal architecture [60], our analyses revealed no strong correlations between either NFκB or YAP activation and any morphological parameter (Supplementary Figure 8). This indicates that the differences in the expression of NFκB and nuclear translocation are driven primarily by scaffold integration, rather than changes in cell shape.

Together, these results demonstrate that collagen + MAP matrices limit fibroblast transition across multiple functional axes, contractility, collagen expression, mechanotransduction, and inflammatory state, while hydrogels promote a pro-fibrotic phenotype. These results collectively show that this effect persists regardless of TGF-β1 stimulation and correlates with the physical integration established between MAP scaffolds and the surrounding ECM.

Distinct fibroblast phenotypes emerge in response to MAP or hydrogel matrix environments.

To determine whether the MAP scaffold modulates fibroblast behavior at the level of cell-state heterogeneity, we performed high-dimensional flow cytometry followed by UMAP embedding and FlowSOM clustering (Figure 8A–C). This analysis enabled simultaneous evaluation of cytoskeletal activation (αSMA) [46], ECM production (Collagen I)[46, 54], fibroblast identity and remodeling potential (CD90) [8, 61–64], mechanotransduction (YAP) [57], and inflamed state (NFκB) [56]. Because fibroblasts adopt a spectrum of morphological and functional phenotypes ranging from quiescent to inflammatory states, we summarized these well-characterized phenotypes in Supplementary Figure 9 to provide a reference framework for interpreting the seven clusters identified in our dataset. Briefly, quiescent fibroblasts exhibit weak cytoskeletal organization, low αSMA and Collagen I expression, diminished YAP and NFκB activity, and variable CD90 levels [46]. Early-activated fibroblasts demonstrate moderate αSMA and CD90 expression [46]. Late transitory fibroblasts or protomyofibroblasts show high expression of collagen [46], pro-fibrotic/myofibroblast phenotypes are defined by strong stress fibers, dense pericellular collagen, and elevated αSMA with reduced nuclear YAP [46, 48, 52]. Transitory myofibroblasts display similar marker expression of pro-fibrotic/myofibroblast phenotypes accompanied by higher levels of YAP [65]. Inflammatory fibroblasts and inflammatory-myofibroblast states exhibit high NFκB activity, prominent stress fibers, and limited CD90 expression [50, 52, 66, 67]. Using this reference, we assigned biological meaning to each FlowSOM-derived subpopulation based on its characteristic marker profile.

**Figure 8.**
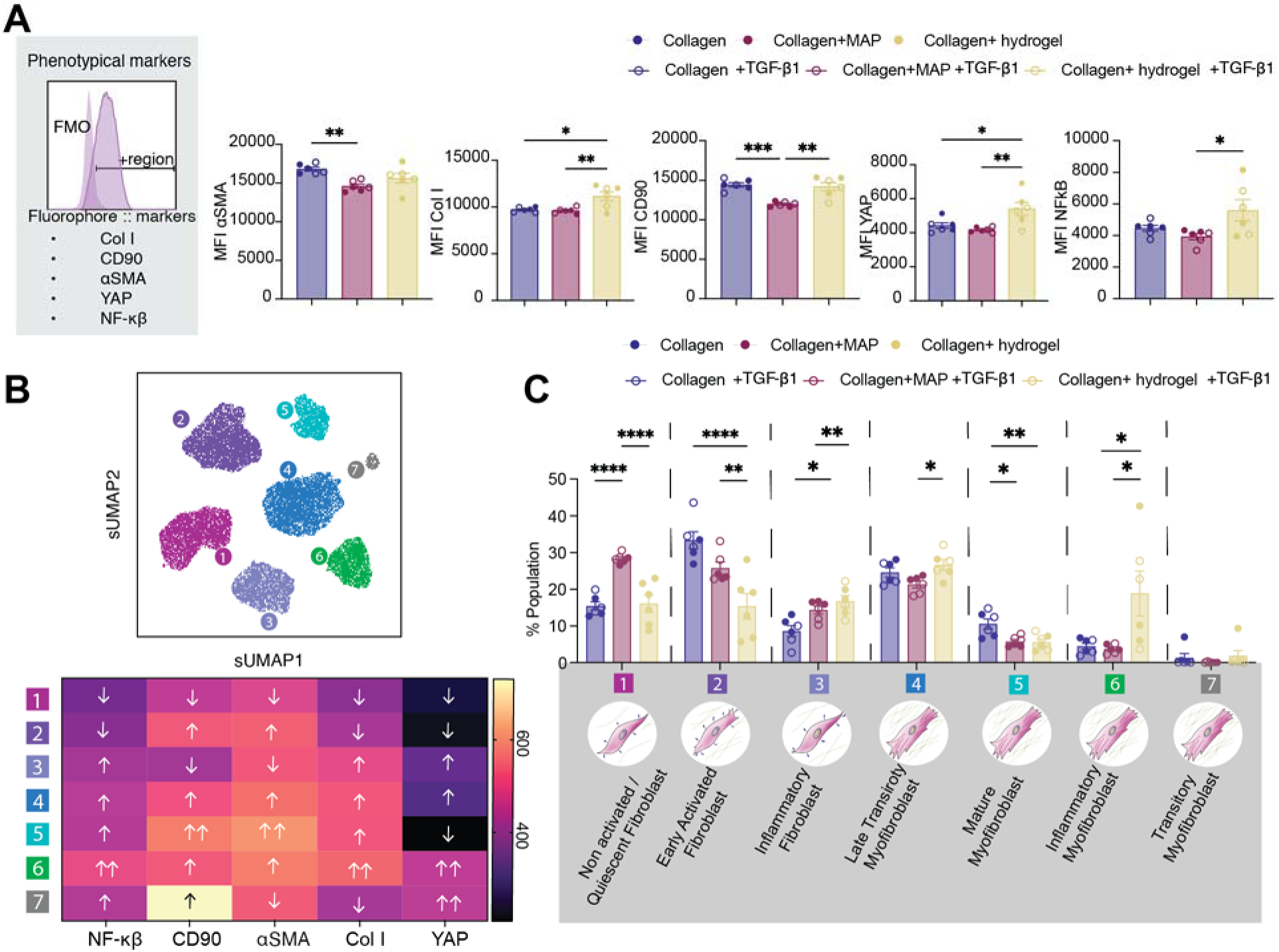
Distinct fibroblast phenotypes emerge in response to MAP or hydrogel matrix environments. (A) Flow cytometry analysis of fibroblast populations from collagen-only, collagen + MAP, and collagen + hydrogel gels with or without TGF-β1 stimulation. Plots show mean fluorescence intensity (MFI) for αSMA, collagen I, CD90, YAP, and NF-κB, demonstrating differences in activation and phenotype based on biomaterial condition. (B) UMAP clustering visualizes heterogeneity in fibroblast phenotypes with seven subpopulations defined by distinct marker expression patterns (heatmap; up = white, down = purple) and functional attributes. (C) Quantification of population percentage for each subcluster under all biomaterial conditions, revealing shifts toward inflammatory, myofibroblast, or quiescent fibroblast subtypes. Data are presented as mean ± S.E.M. Filled circles represent measurements from samples without TGF-β1, open circles correspond to samples with TGF-β1. Statistical analysis was performed using one-way ANOVA with Tukey’s post-hoc test (*p<0.05, **p<0.01, ***p<0.001, ****p<0.0001). N = Number of collagen gels, each with independent cell cultures.

Comparable to the immunofluorescence findings, TGF-β1 stimulation did not substantially alter the distribution of fibroblast subpopulations. Across all conditions, TGF-β1 produced only modest changes that did not reorganize cluster structure, confirming that the dominant cues regulating fibroblast state were mechanical rather than biochemical in this system. This observation agrees with reports that fibroblasts in prestressed collagen matrices can maintain activation independent of exogenous TGF-β1 [37, 53] and aligns with the minimal TGF-β1–dependent effects observed above (Supplementary Figure 5).

Across matrix conditions, seven distinct fibroblast states emerged, exhibiting combinations of quiescent, mechanically primed, ECM-producing, myofibroblast-like, and inflammatory phenotypes. Notably, collagen + MAP matrices were enriched for a quiescent subpopulation (Population 1) characterized by uniformly low αSMA, Collagen I, CD90, YAP, and NFκB expression. This state, which aligns with the non-activated/ quiescent phenotype, accounted for (∼30%) in collagen + MAP matrices compared with collagen-only (∼10–20%) or collagen + hydrogel matrices (<10%) (Figure 8C). The enrichment of this quiescent population in MAP scaffolds is consistent with the reduced compaction and suppressed marker expression observed above. In contrast, collagen + hydrogel matrices displayed a markedly different distribution of phenotypes. Subpopulations characterized by elevated YAP, αSMA, Collagen I, and NFκB, corresponding to the mechanically primed, activated myofibroblast, and inflammatory myofibroblast phenotypes (Supplementary Figure 9), were significantly enriched in the hydrogel condition.

Fibroblasts in collagen-only matrices were distributed across intermediate phenotypes, including later transitory fibroblasts expressing high αSMA but low Collagen I, and populations displaying moderate activation across all markers. These intermediate states mirror the partial compaction and midrange activation marker expression observed in collagen-only gels and reinforce the view that scaffold architecture and the resulting mechanical environment directly shape fibroblast identity. Analysis of proliferation and apoptosis are more associated with cell state rather than with material type (Supplementary Figure 10). Inflammatory and activated myofibroblast populations exhibited the highest apoptotic fractions, whereas quiescent fibroblasts showed the lowest, indicating that differences among biomaterial conditions reflect shifts in fibroblast phenotype rather than changes in total cell number.

Together, these results demonstrate that MAP scaffolds alter the distribution of fibroblast phenotypes by enriching quiescent phenotypes and suppressing both late transitory and inflammatory myofibroblast phenotypes compared with collagen + hydrogel conditions. In contrast, hydrogels promote contractile, ECM-producing, and NFκB-activated fibroblast states associated with fibrotic remodeling. These findings establish scaffold architecture as a key regulator of fibroblast heterogeneity and highlight MAP scaffolds as potent suppresses of both mechanical and inflammatory aspects of fibroblast phenotype.

## Discussion

The goal of this study was to determine how the physical integration of biomaterials with the extracellular matrix (ECM) influences fibroblast phenotype. While MAP scaffolds have been widely recognized for their regenerative capacity in vivo, the mechanisms underlying their anti-fibrotic effects have remained unclear. Here, using a reductionist, in vitro model, we demonstrate that the granular architecture of MAP scaffolds enables seamless integration with collagen type I matrices, thereby significantly promoting ECM-biomaterial physical integration and preventing fibroblast activation. Together, these findings establish physical matrix–biomaterial integration as a key determinant that controls fibroblast activation.

Despite sharing identical chemistry and similar bulk mechanical properties, MAP scaffolds and traditional hydrogels interact with collagen in strikingly different ways. Collagen fibers are assembled within the void spaces of MAP scaffolds, creating a continuous composite material. In contrast, hydrogels formed a continuous gap that acted as a slip plane and led to material detachment under mechanical load. In the cellular system, this lack of integration led to high compaction and distortion of collagen architecture. These observations highlight that architectural differences at the biomaterial interface produce distinct mechanical microenvironments that ultimately shape fibroblast phenotype.

A central finding of this work is the suppression of NF-κB expression and nuclear localization in fibroblasts within collagen + MAP matrices. NF-κB is a master regulator of inflammatory and fibrotic responses, integrating biochemical cues with mechanical stress [68–70]. Across imaging, compaction assays, and flow cytometry, MAP scaffolds consistently reduced NF-κB expression and nuclear localization, even in the presence of TGF-β1. The emergence of a dominant quiescent fibroblast population in MAP-containing matrices, together with reduced inflammatory and mechanically associated markers, indicates that physically integrated environments suppress both contractile and inflammatory markers associated with fibroblast activation. In contrast, collagen + hydrogel matrices enriched for fibroblast states characterized by high αSMA, Collagen I, YAP, and NF-κB, indicating that physical discontinuities can promote a more pro-fibrotic and inflammatory phenotype.

Because cell number, proliferation, and apoptosis did not explain these differences, the results point to physical continuity as the dominant regulator of fibroblast phenotype. The physical continuity imparted by MAP scaffolds eliminates sharp interfacial discontinuities and prevents large-scale collagen distortion, whereas slip-plane formation in hydrogel matrices generates core detachment and increase in collagen compaction. This distinction provides a physical framework through which mechanosensitive cellular responses, including those associated with inflammatory activation, may be differentially influenced in MAP versus hydrogel systems.

These findings motivate important future directions. A key next step will be to determine how the local matrix mechanics and connectivity created by MAP–ECM integration influences NF-κB activation, YAP responses, and fibroblast transition. Future experiments will directly investigate the proposed mechanisms by examining the upstream and downstream pathways that drive changes in NF-κB expression and fibroblast phenotype. Potential mechanisms include the following. Mechanically induced ECM compaction may expose or release endogenous danger-associated molecular patterns (DAMPs) [71], including fibronectin-EDA, tenascin-C, HMGB1, or fragmented collagen [72]., which can activate TLR4 signaling and drive MyD88-dependent IKK activation and NF-κB nuclear translocation [73, 74]. ECM compaction and remodeling may also promote the release of latent matrix-bound TGF-β, reinforcing NF-κB activation through established TGF-β–NF-κB crosstalk [75, 76]. In parallel, matrix mechanics may directly regulate NF-κB through mechanotransduction pathways, including integrin engagement, cytoskeletal tension, and mechanosensitive ion channels [77, 78]. Secondary inflammatory signaling, such as TNF-α or IL-1β receptor activation, may further amplify NF-κB activity downstream of these mechanically initiated events [56]. Moreover, systematic variation of MAP scaffold design parameters (e.g., microgel stiffness, packing density, and pore size) will help determine how specific architectural features tune physical continuity and thereby regulate fibroblast phenotype. Such studies will clarify whether granular scaffolds can be rationally engineered to modulate fibroblast activation through precisely controlled mechanical integration with the ECM.

In summary, this work demonstrates that integration of granular MAP scaffolds with collagen ECM modulates fibroblast phenotype by reducing NF-κB expression and nuclear localization, and reorganizing ECM architecture. These results position physical integration as a powerful design principle for creating biomaterials that resist fibrosis and support regenerative healing.

## Methodology

### Collagen isolation

Collagen was isolated from rat tails (Charles River, 8-9 weeks old) following a modified version of the protocol described by Rajan et al[79]. Collagen fibers were extracted from rat tails and sequentially washed with acetone, 70% (v/v) isopropanol, and 0.02 N acetic acid. The cleaned fibers were then dissolved in 0.5 N acetic acid and stirred at 4°C for a minimum of 72 hours to ensure complete solubilization. The resulting collagen solution was filtered through a 23G gauge needle and centrifuged at 14,000 x g for 45 minutes. The supernatant was carefully collected and dialyzed against three changes of 0.02 M acetic acid at 4°C, over a three-day period. The final transparent solution (∼ 5 mg/mL) of concentrated collagen was stored at 4°C until use.

### Collagen labelling

To study collagen architecture, collagen fibers were fluorescently stained based on the protocol by Doyle [80]. First, a collagen gel of 2 mg/mL was prepared from a stock concentration of 5 mg/mL using 10X DMEM and 10X reconstitution buffer for osmolarity balance. Then, 1 N NaOH was added until reaching a pH of 7. The collagen was allowed to polymerize for 1 hour. Once the gel was completely polymerized, it was incubated in a borate buffer solution (pH 9) for 15 minutes at room temperature. The collagen was then labeled using a solution of Atto-488 NHS-ester in borate buffer and incubated for 1 hour at room temperature. The gel was washed with Tris-buffer (pH 7.5) and PBS at least 6 times over 4 hours to remove excess dye. The collagen solution was then liquefied using 500 mM acetic acid and gentle stirring until the sample was completely liquid. The labeled collagen was subsequently dialyzed against 20 mM acetic acid. Finally, the collagen was aliquoted and stored at 4°C, protected from light until use.

### Material fabrication

Both the MAP scaffold and hydrogel were fabricated using PEG-based gels that incorporate cleavable crosslinker peptides, in line with previous publications. To prepare the precursor solution, 10% (w/v) 8-arm PEG-Vinyl sulfone (PEG-VS) 20 kDa (Jemkem) was dissolved in a buffer containing 300 mM triethanolamine (TEOA) at pH 8.25. Subsequently, 1000 µM of RGD peptide (Ac-RGDSPGERCG-NH2, Genscript) was added and incubated for 45 minutes at 37°C. The pH of the solution was adjusted to 4.5 using 5 N HCl. To create the gels, the PEG-VS solution was mixed in a 1:1 ratio with a solution containing 12 mM of L-MMP peptide (Ac-GCRDGPQGIWGQDRCG-NH2, Genscript) and 5 mM TCEP in Milli-Q water.

Microgels for the MAP scaffold were produced using an oil-based microfluidic setup, following the protocol by Griffin et al. Briefly, heavy mineral oil with 5% Span80 was utilized to form microgels by pinching the precursor solution. The microgels were solidified through pH-mediated chemistry involving heavy mineral oil with 5% Span80 and 2% Trimethylamine (TEA) in the collection tube. The microgels were then subjected to multiple washes using buffer exchange through centrifugation with 0.3 M HEPES, both with and without 2% Pluronic. For the final wash, microgels were added to a solution of 0.3 M HEPES containing 10 mM CaCl_2_.

For norbornene annealing, the microgels were post-conjugated using a solution of 10 µM Methyltetrazine-PEG4-Maleimide (Click Chemistry Tools) (Supplementary Figure 1). After conjugation, the microgels were washed and stored at 4°C in either 0.3 M HEPES or culture media until needed. For the annealing process, a solution of 9% (w/v) 8-arm PEG-NB 20 kDa (JemKem) was employed.

For hydrogel preparation, the precursor solution was gelled using 3.4% (v/v) of 5 M NaOH. Immediately thereafter, the solution was placed between two Sigma-coated cover glasses, with Teflon spacers of 1 mm thickness, for 10 minutes. The gels were stored at 4°C in either 0.3 M HEPES or culture media until use.

To label the material, 20 µM of Alexa Fluor 647 Maleimide was incorporated into the precursor solution prior to gelation.

### Acellular model of collagen matrices

To investigate the interaction between MAP scaffolds or hydrogels and collagen fiber structure and formation, both materials were embedded into collagen matrices. A volume of 10 µL of fluorescently labeled MAP scaffold or hydrogel was added to the center of a PDMS molds[81]. The MAP scaffolds were incubated for 45 minutes at 37°C to allow annealing. Subsequently, the molds were pre-chilled at 4°C, and 25 µL of 2 mg/mL collagen gel containing 0.2% labeled collagen (prepared following the protocol described by Doyle[80, 82]) was added. To ensure that the collagen filled the entire mold, the collagen matrices were centrifuged at 100×g for 3 minutes at 4°C. The samples were then incubated overnight at 4°C to allow proper gelation (Figure 2a). After the collagen matrices had gelled, 100 µL of 1X PBS was added to the molds, and the samples were imaged using a Nikon C2+ point-scanning confocal microscope. Images were collected at either 20× or 40× magnification with a resolution of 1024 pixels and a Z-stack interval of 0.5 µm to achieve sufficient resolution for rendering collagen fibers.

### Collagen fibers analysis

The Filament Tracer tool in Imaris was used to analyze the architecture of collagen fibers. Initially, images were converted into .ims format, and a surface was created to mask the collagen fibers, effectively removing background noise. The tool was then applied with a specified fiber diameter range of 0.4 µm to 6 µm, selecting the option for “fibers without spine.” To examine regions at varying distances from the biomaterial, three regions of interest (ROIs) were chosen per image, each with voxel dimensions of 50 µm × 50 µm × 50 µm. Region R1 corresponded to areas within <50 µm of the biomaterial (MAP or hydrogel). Region R2 encompassed areas approximately 50–100 µm from the biomaterial. Region R3 included areas located at >100 µm from the biomaterial.

Since each image was rendered independently, all collected data, such as area, volume, orientation, and diameter, were normalized internally by averaging each property within its respective image. This approach ensured consistency and comparability across samples.

### Cellular model: Modified contraction assay

A modified contraction assay was employed to evaluate the interaction of MAP or hydrogel within cellular collagen matrices. First, collagen solutions (2 mg/mL) containing 0.1% labeled collagen were prepared and mixed with NIH/3T3 cell pellets to achieve a final concentration of 3×10L cells per mL of collagen. 200 µL of this cellular collagen were transferred to a 48-well plate. Immediately, 10 µL of MAP or hydrogel was added to the center of each matrix, followed by incubation at room temperature until complete collagen gelation. Subsequently, 500 µL of DMEM supplemented with 1% antibiotic-antimycotic (AA) was added to each well, and the samples were incubated at 37°C for 48 hours. After this period, matrices were carefully released from the wells using a sterile plastic spatula (stressed matrix model[37]), and collagen contraction rate was quantified over the next five hours. To activate the cells, 10 ng/mL of TGF-β1 was added to designated wells during incubation. For assay validation, 10 mM BDM (2,3-butanedione monoxime) was included as a negative control to confirm the absence of contraction (Supplementary Figure 5), ensuring reproducibility and robustness of the experimental setup.

### Flow cytometry

After contraction, cells embedded in the collagen matrices were isolated using mechanical disruption with a displacement pipette in digestion buffer containing Liberase (RPMI, 5% FBS, DNase I at 50 U/mL, and 300 µg/mL Liberase [Roche]). The mixture was incubated for 10 minutes at 37°C. The isolated cells were washed, pellet down and labeled for viability using Zombie NIR (BioLegend) for 15 minutes at room temperature (RT). Subsequently, cells were blocked using FcR Blocking Buffer (Miltenyi Biotec) for 10 minutes at 4°C. Extracellular markers (Supplementary Table 1) were then stained for 30 minutes at 4°C. After staining, the cells were fixed and permeabilized using the Foxp3/Transcription Factor Staining Buffer Set (eBioscience™). Following fixation and permeabilization, the cells were washed and stained for intracellular markers (Supplementary Table 1) for an additional 30 minutes at 4°C. Finally, the cells were washed and analyzed using a Cytek Northern Lights Spectra cytometer.

The analysis of the resulting .fcs files was performed using FlowJo V10 software, with plugins such as FlowClean, DownSample, UMAP, and FlowSOM[83] for data processing.

### Immunofluorescence

After contraction, multiple sets of cellular collagen matrices were fixed with 4% paraformaldehyde (PFA) in HBSS buffer (pH 7.4) for 30 minutes at room temperature (RT). The matrices were then washed with PBS and incubated for 2 hours at RT in 1X PBS containing 1% Triton X-100 and 3% BSA. Following this, the cells were incubated overnight at RT with diluted primary antibodies prepared in a solution of 0.3% Triton X-100 and 3% BSA (Supplementary Table 2). After antibody incubation, the gels were stained with DAPI for 5 minutes and subsequently washed multiple times with 0.3% Triton X-100 in PBS. A final wash was performed using 1X PBS, and the matrices were kept hydrated in 500 µL of 1X PBS until imaging. Images were acquired using a Nikon C2+ point-scanning confocal microscope at 10X and 60X magnification, with a resolution of 1024 pixels. Image analysis and quantification of marker intensity, protein localization, collagen intensity and coherence, as well as cell morphology, were performed using ImageJ/Fiji software.

To calculate the Chromatin Condensation Parameter (CCP), we used ImageJ/Fiji and followed the previously described protocol[84]. Briefly, we duplicated the regions of interest (ROI) of the nuclei from the original 60X images. We applied the “Find Edges” function for edge detection, thresholded and binarized the image, and calculated the total edge ratio by dividing the area of the edges by the total nuclear area.

### Imagining analysis of MAP scaffolds using LOVAMAP

A volume of 10 µL of fluorescently labeled MAP scaffold was added to PDMS molds [85]. After 45-60 minutes of incubation at 37°C in a humidified chamber, the microgel was completely annealed. Then, 100 µL of CUBIC-Reagent II [86] was added to the molds, and the samples were imaged using a Nikon C2+ point-scanning confocal microscope. Images were collected at 20x magnification with a resolution of 1024 pixels and a Z-stack interval of 0.25 µm to achieve sufficient resolution for rendering the individual microgels. The collected images were then formatted for LOVAMAP input by running custom code to voxelize and segment individual microgels [34]. LOVAMAP analyzes the MAP scaffolds by segmenting the void space into 3D pores (i.e., open pockets of space). We reported seven pore metrics previously described in Riley et al. Briefly, longest length (µm) is the longest end-to-end length of a pore. Shortest length (µm) is the average width along the internal backbone of the pore. Largest door (µm) is the width of the largest opening (“door”) into or out of the pore along a backbone, while smallest is as the name suggests. Volume (pL) of a pore is reported in picoliters, and surface area (µm^2^/1000) is scaled by 1000 to match the length scale of the reported volume. Isotropy is a unitless measurement that ranges between 0 and 1, where a score of 0 indicates pore orientation that is orthogonal to the average orientation and a score of 1 indicates pore orientation that is parallel to the average [33].

### Statistical analysis

All experiments were conducted in triplicate using independent batches of collagen gels and cells from separate cultures. Statistical analysis was performed using GraphPad Prism software 10 (GraphPad Software). One-way ANOVA followed by Tukey’s post hoc test was employed to compare the effects of material treatments (MAP, Hydrogel, or collagen alone), with p-values less than 0.05 considered statistically significant. For specific datasets, two-way ANOVA was utilized to assess the significance of both material treatment and activation (with and without TGF-β1). Single collagen fiber analyses were evaluated using nested ANOVA with Dunnett’s post hoc multiple comparison test.

## Acknowledgements

We would like to thank the National Institutes of Health (R01AI152568) and all the members of Segura at Duke University for their support. Thank you to Duke University Light Microscopy Core Facility (LMCF) and their staff, and Amber English, Applications Scientist at Imaris, An Oxford Instruments Brand for their support for the analysis of collagen fiber architecture.

## Author contributions

ASA made substantial contributions to the design of the work, acquisition, analysis, and interpretation of data. MH performed part of the experiment that contributed substantially to Figure 6 and Supplementary Figures 4 to 6. CR performed part of the experiment that contributed substantially to Figure 5. AK performed part of the experiment that contributed substantially to Figure 1. LZ performed part of the experiment that contributed substantially to Supplementary Figure 1 and 2. LR provided conceptualization, data, and analysis that contributed substantially to Figure 1 and Supplementary Figure 2. ASA drafted the manuscript, and all the authors discussed the results and contributed to writing portions of the manuscript and editing the manuscript. BDH and TS provided guidance and discussion throughout the project, and made substantial contributions to experimental design, data analysis, and manuscript editing.

## Supplementary Figures

**Supplementary Figure 1.**
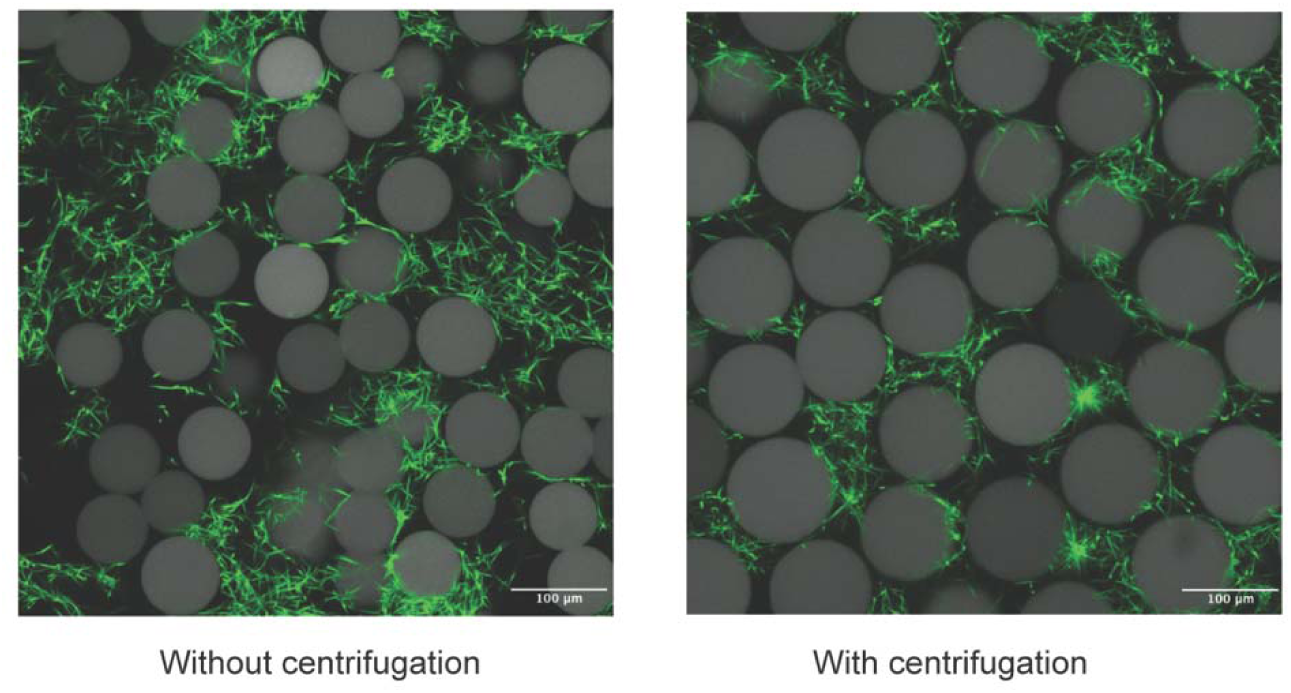
Centrifugation promotes homogeneous collagen infiltration within MAP scaffolds in the acellular model. Representative confocal images showing collagen fibrillogenesis (green) within microporous annealed particle (MAP) scaffolds under two conditions: without centrifugation and with centrifugation during collagen polymerization. In the absence of centrifugation, collagen fibers distribute unevenly throughout the scaffold pores, resulting in regions of sparse fibrillar network formation. In contrast, centrifugation facilitates uniform infiltration of the collagen solution throughout the interstitial spaces of the MAP scaffold prior to fibrillogenesis, producing a more homogeneous collagen fiber network across the scaffold pores. Scale bars: 100 μm.

**Supplementary Figure 2.**
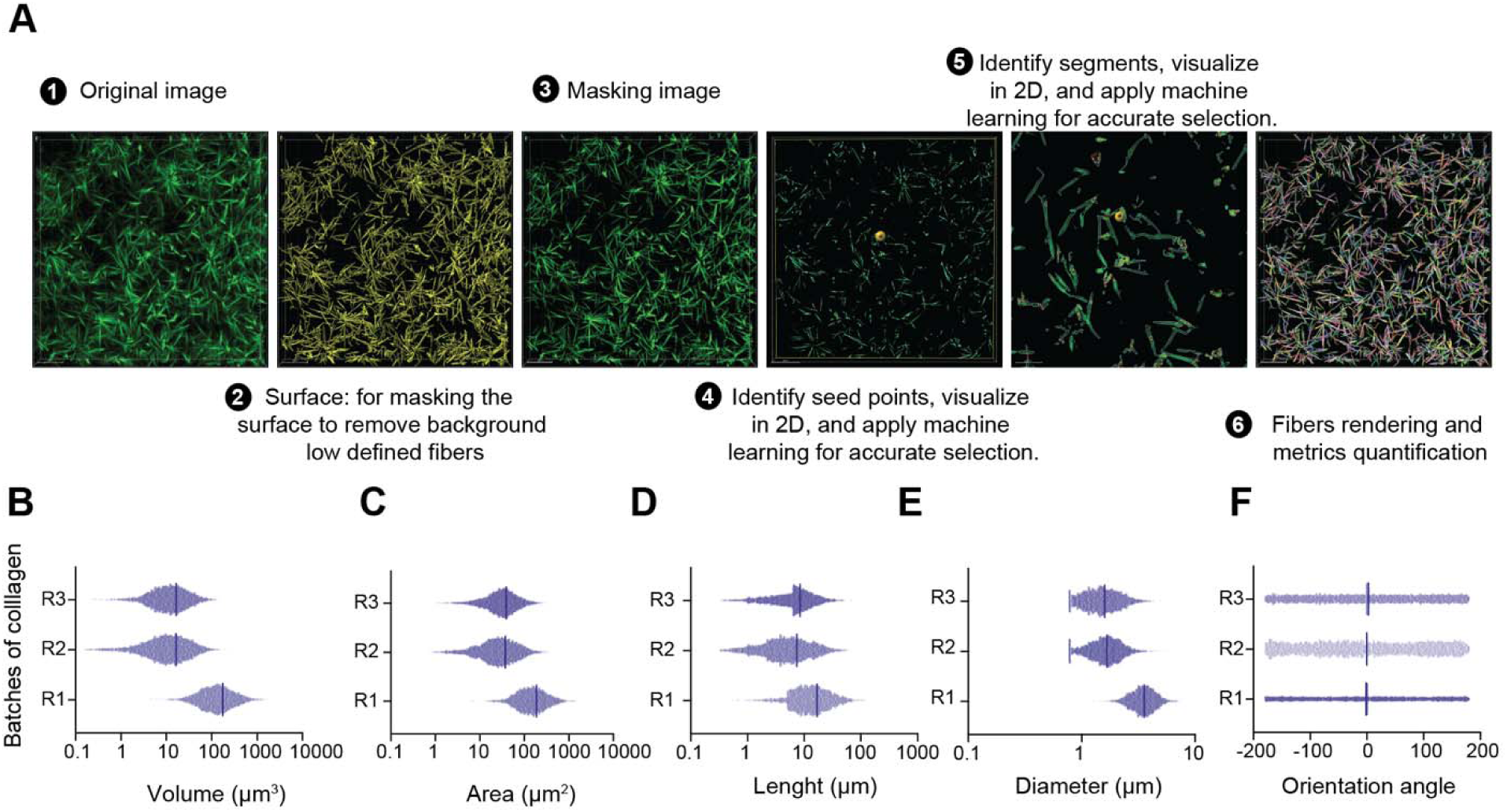
Quantification of collagen fibers using IMARIS. (A) Workflow for extracting and analyzing collagen fiber metrics: (1) original confocal image, (2) surface masking to eliminate poorly defined fibers and background, (3) refined masking, (4–5) identification and segmentation of fibers using Filament tracer tool for accurate selection, and (6) rendering for quantification of architectural metrics. (B–F) Distribution of collagen fiber volume (B), area (C), length (D), diameter (E), and orientation angle (F) for three independent collagen batches (R1, R2, R3), demonstrating reproducibility and structural variability among experimental replicates.

**Supplementary Figure 3.**
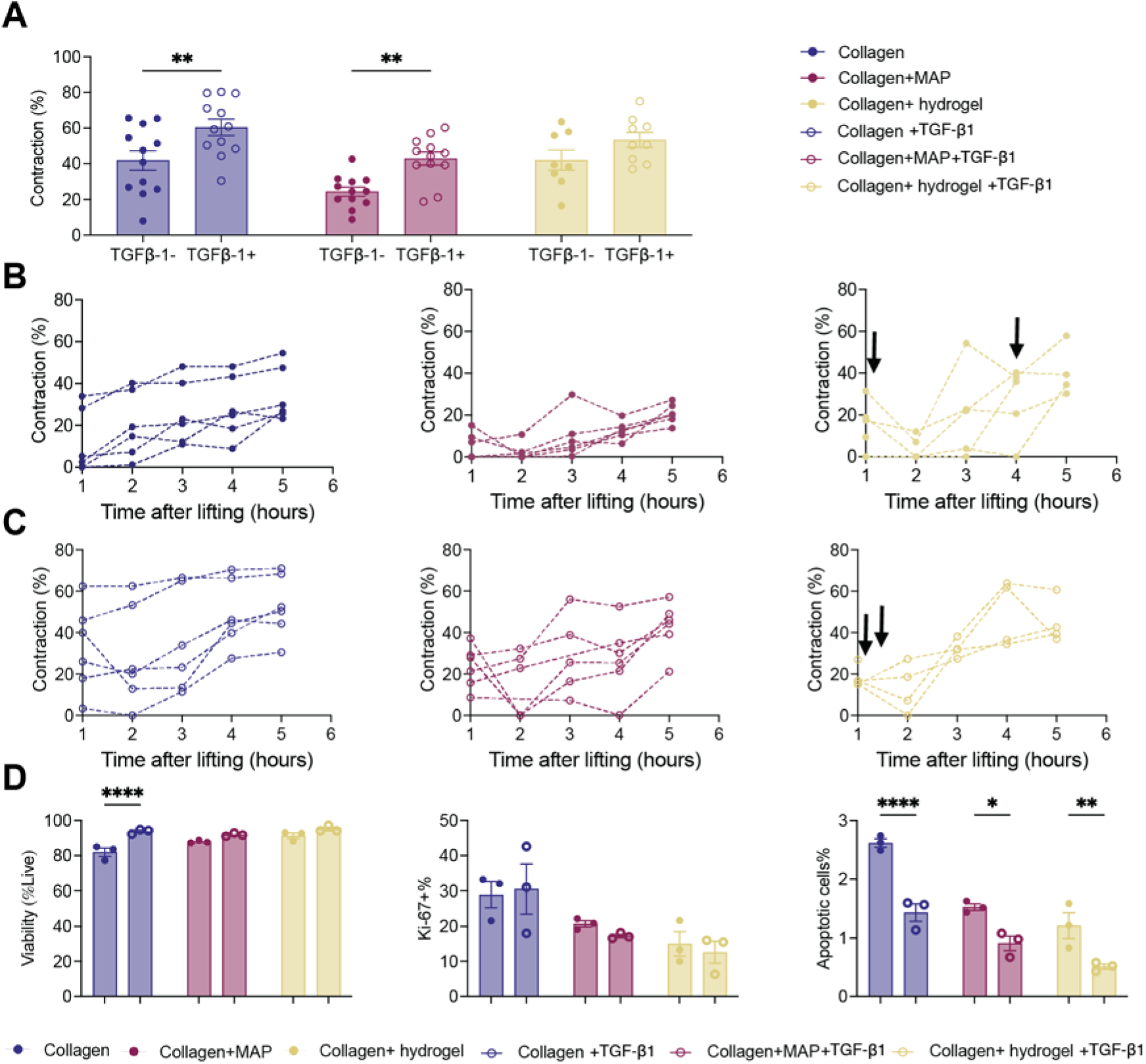
Time course of collagen gel compaction for each collagen matrix. (A) Quantification of compaction reveals that TGF-β1 increases compaction compared to collagen matrices without TGF-β1, regardless of biomaterial presence. (B) Compaction curves for collagen-only, collagen + MAP, and collagen + hydrogel systems without TGF-β1; (C) Curves for the same conditions with TGF-β1. Arrows in the hydrogel curves indicate the time points when the hydrogel core detached from the surrounding matrix, resulting in a sharp increase or change in contraction. (D) Comparative analysis of fibroblast viability, apoptosis, and proliferation (Ki-67+) across collagen matrix types of fibroblasts activated with TGF-β1 or without TGF-β1. Each line represents a separate experimental replicate for the indicated condition. Data are presented as mean ± S.E.M. Filled circles represent measurements from samples treated without TGF-β1, while open circles correspond to samples treated with TGF-β1. Statistical analysis was performed using two-way ANOVA with Tukey’s multiple comparison test (*p<0.05,**p<0.01, ***p<0.001, ****p<0.0001) N = Number of collagen gels; each point represents an independent cell culture replicate.

**Supplementary Figure 4.**
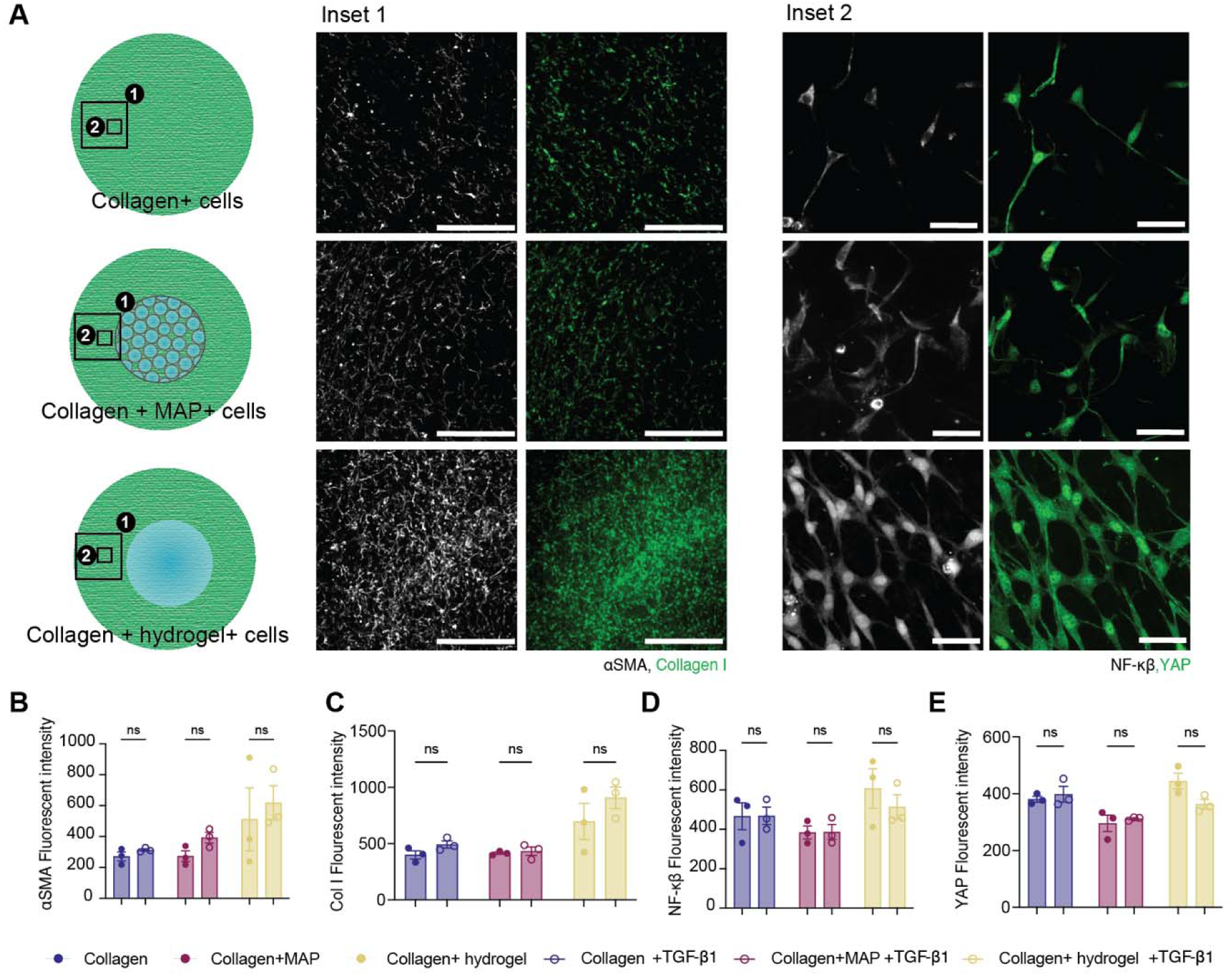
Fibroblast activation in collagen matrices with MAP or hydrogel occurs independently of TGFβ1. (A) Representative fluorescence images from collagen-only, collagen + MAP, and collagen + hydrogel constructs showing marker localization for Inset 1 (20X): αSMA (white) and collagen I, and Inset 2 (60X): (green), NF-κB (white), and YAP (green). Schematics indicate the analyzed regions for each condition. Panels show both tissue-level and cellular insets. (B–E) Quantification of αSMA (B), collagen I (C), NF-κB (D), and YAP (E). Data are displayed as mean ± S.E.M. and individual values. Filled circles represent measurements from samples treated without TGF-β1, while open circles correspond to samples treated with TGF-β1. Statistical analysis was performed using one-way ANOVA with Tukey’s multiple comparison test (ns: not significant). N = Number of collagen gels; each point represents an independent cell culture replicate.

**Supplementary Figure 5.**
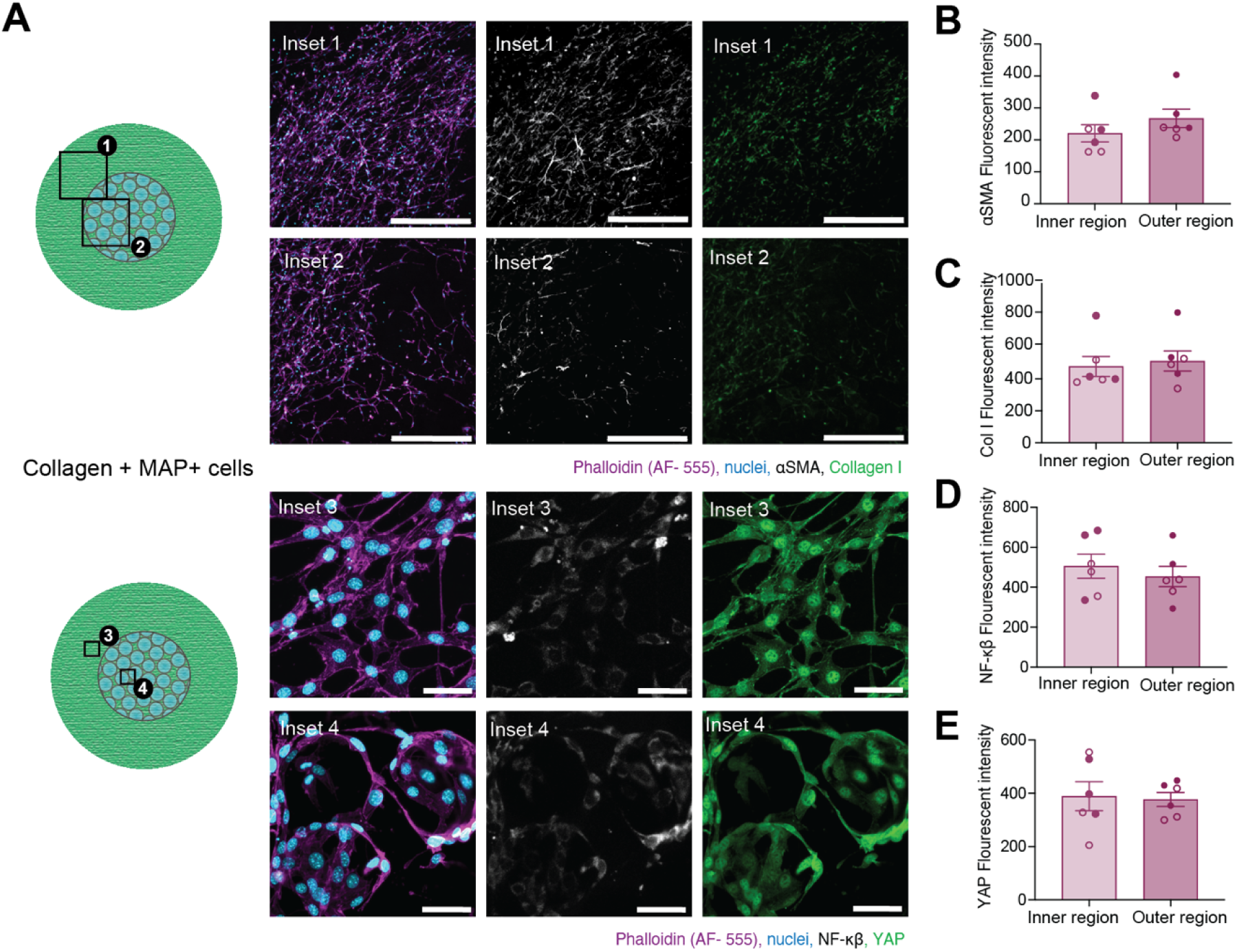
Expression of fibroblast activation markers is consistent across inner and outer regions of Collagen + MAP gels. (A) Representative confocal fluorescence images from inner and outer regions of collagen + MAP gels, stained for actin filaments (phalloidin, magenta), nuclei (blue), αSMA (white), collagen I (green), NF-κB (white), and YAP (green). Schematics indicate sampling regions within the gels. Insets 1 and 2 (20X) and Insets 3, and 4 (60X). (B–E) Quantification of αSMA (B), collagen I (C), NF-κB (D), and YAP (E) fluorescent intensity reveals no significant differences between inner and outer gel regions. Data are displayed as mean ± S.E.M. and individual values. Filled circles represent measurements from samples treated without TGF-β1, while open circles correspond to samples treated with TGF-β1. Statistical analysis was performed using one-way ANOVA with Tukey’s multiple comparison test (ns: not significant). N = Number of collagen gels; each point represents an independent cell culture replicate.

**Supplementary Figure 6.**
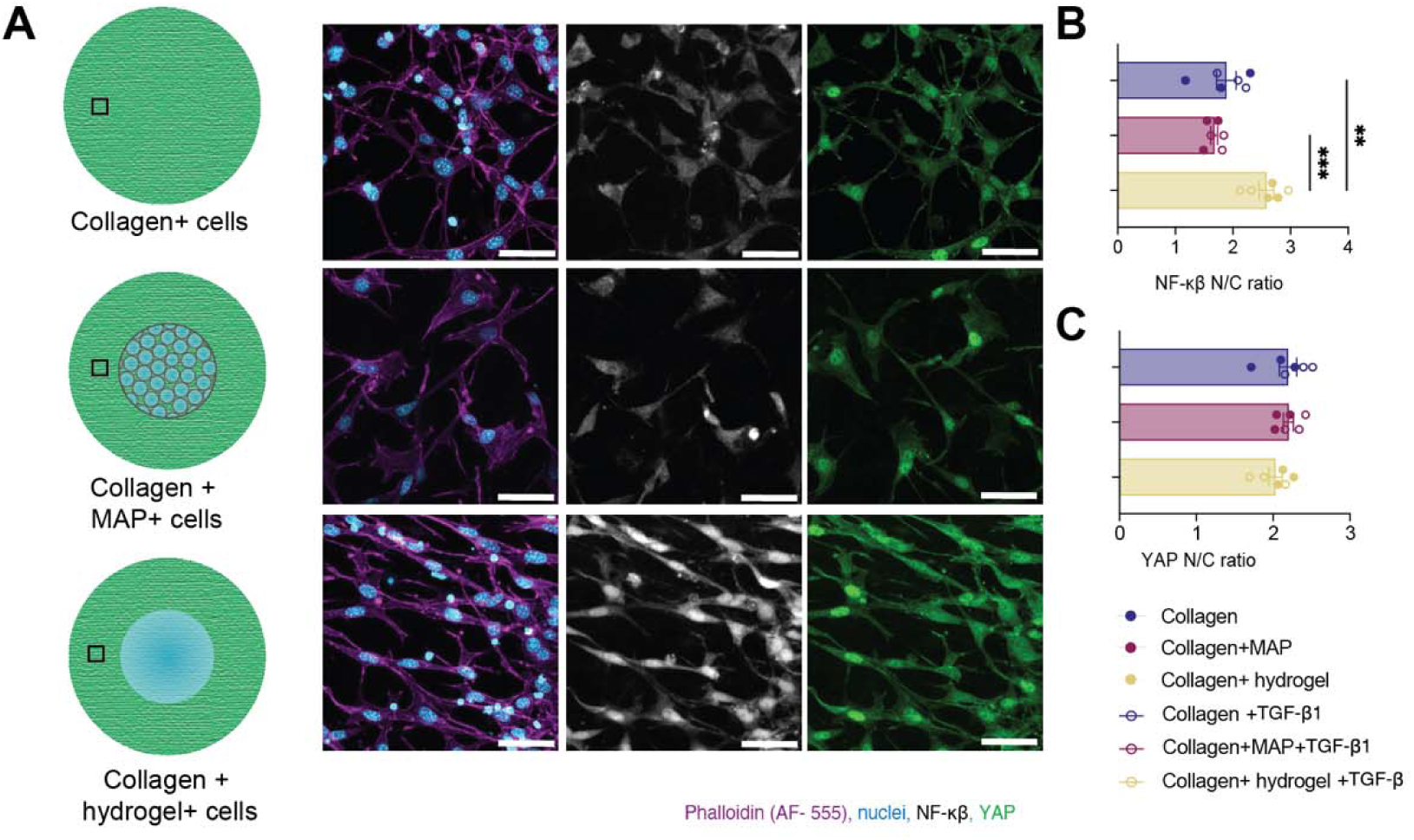
Nuclear NF-κB activation is elevated in fibroblasts within collagen + hydrogel matrices. (A) Representative confocal fluorescence images of collagen-only, collagen + MAP, and collagen + hydrogel gels, showing actin filaments (phalloidin, magenta), nuclei (blue), NF-κB (white), and YAP (green) staining. Schematics indicate experimental material setup and sampling regions. (B, C) Quantification of NF-κB and YAP Nuclei/cytoplasmic (N/C) ratios for each condition, illustrating material-dependent differences in signaling pathway activity. Data are displayed as mean ± S.E.M. and individual values. Filled circles represent measurements from samples treated without TGF-β1, while open circles correspond to samples treated with TGF-β1. Statistical analysis was performed using one-way ANOVA with Tukey’s multiple comparison test (**p<0.01, ***p<0.001). N = Number of collagen gels; each point represents an independent cell culture replicate.

**Supplementary Figure 7.**
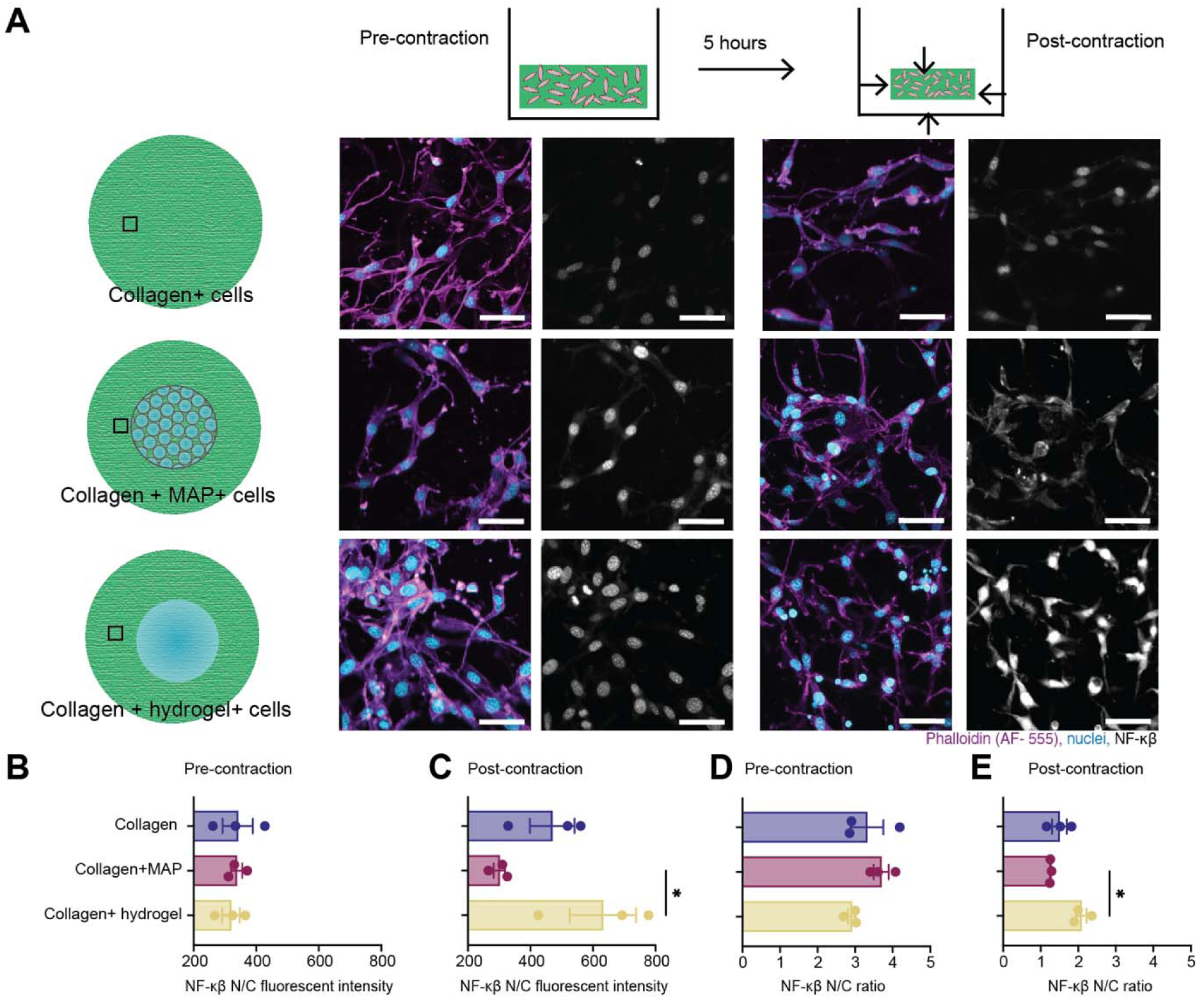
Biomaterial-dependent differences emerge post-compaction. (A) Representative confocal fluorescence images of fibroblasts within collagen-only, collagen + MAP, and collagen + hydrogel constructs, captured before and after gel compaction. Images show actin filaments (phalloidin, magenta), nuclei (blue), and NF-κB (white). Schematic illustrates the experimental workflow and time points. (B,C) Quantification of NF-κB nuclear/cytoplasmic (N/C) fluorescence intensity and N/C ratio pre-contraction; (D,E) Corresponding NF-κB N/C intensity and ratio post-contraction. Data are displayed as mean ± S.E.M. and individual values. Filled circles represent measurements from samples treated without TGF-β1. Statistical analysis was performed using one-way ANOVA with Tukey’s multiple comparison test (*p<0.05). N = Number of collagen gels; each point represents an independent cell culture replicate.

**Supplementary Figure 8.**
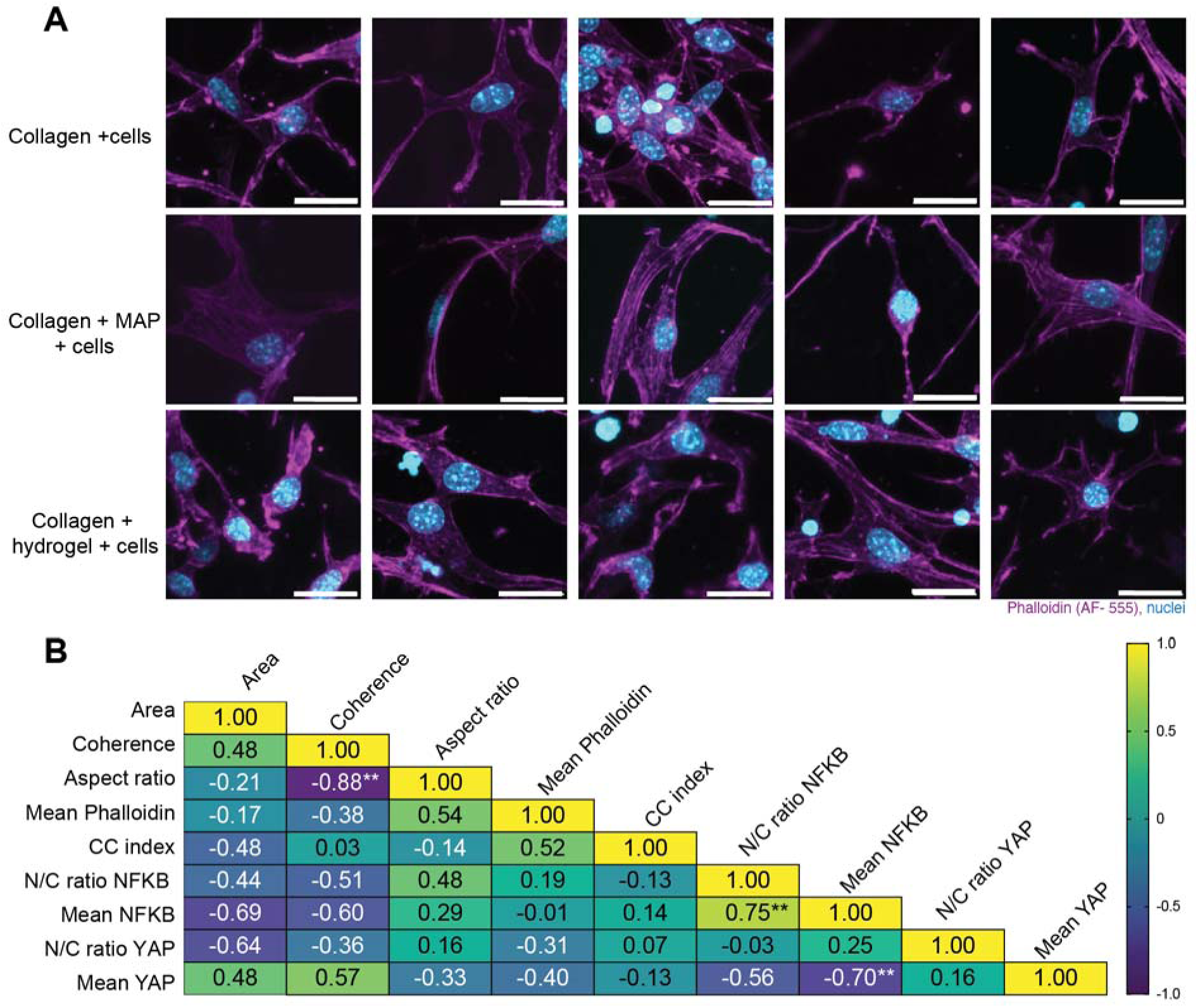
Changes in expression and activation of YAP or NFκB signaling occur independently of fibroblast morphology. (A) Representative single-cell confocal fluorescence images of collagen-only, collagen + MAP, and collagen + hydrogel constructs, stained for F-actin (phalloidin, magenta) and nuclei (blue), highlighting differences in fibroblast morphology and cell shape across material environments. (B) Correlation matrix heatmap quantifying the relationships between cell shape, cytoskeletal, chromatin, and signaling features, including coherence, cell area, aspect ratio, actin intensity, chromatin condensation, and nuclear/cytoplasmic ratios of NF-κB and YAP. Color scale reflects correlation coefficients, with values ranging from -1 (blue) to 1 (yellow. For quantification, 9 cells were analyzed per biological replicate, with 3 biological replicates per collagen gel type. All morphological metrics were paired for the correlation analysis. Pearson correlation test (*p<0.05, **p<0.01,). Scale bars= 20 µm

**Supplementary Figure 9.**
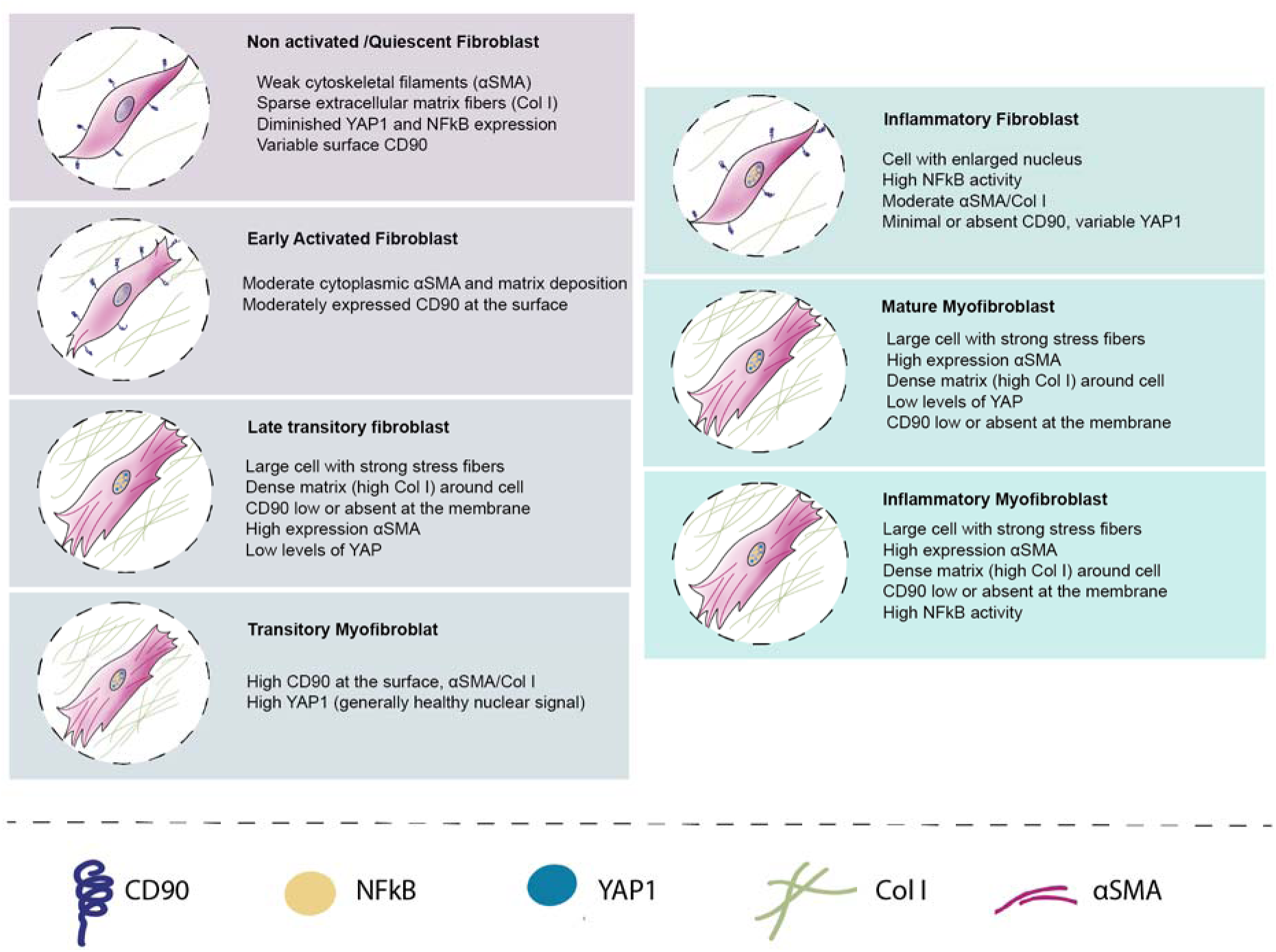
Schematic of fibroblast phenotypes. Distinct morphological and functional states ranging from quiescent to inflammatory fibroblasts, including quiescent, early activated, late transitory fibroblast, transitory myofibroblast, inflammatory fibroblast, and mature and inflammatory myofibroblast types. Each panel depicts characteristic features related to cytoskeletal organization, extracellular matrix composition, and nuclear signaling activity in different fibroblast states. Symbols: Cytoskeletal filaments (αSMA) are shown as magenta lines, collagen type I fibers (Col I) as green fibers, nuclear YAP1 as blue dots, nuclear NFκB as yellow circles, and surface CD90 as dark blue spirals.

**Supplementary Figure 10.**
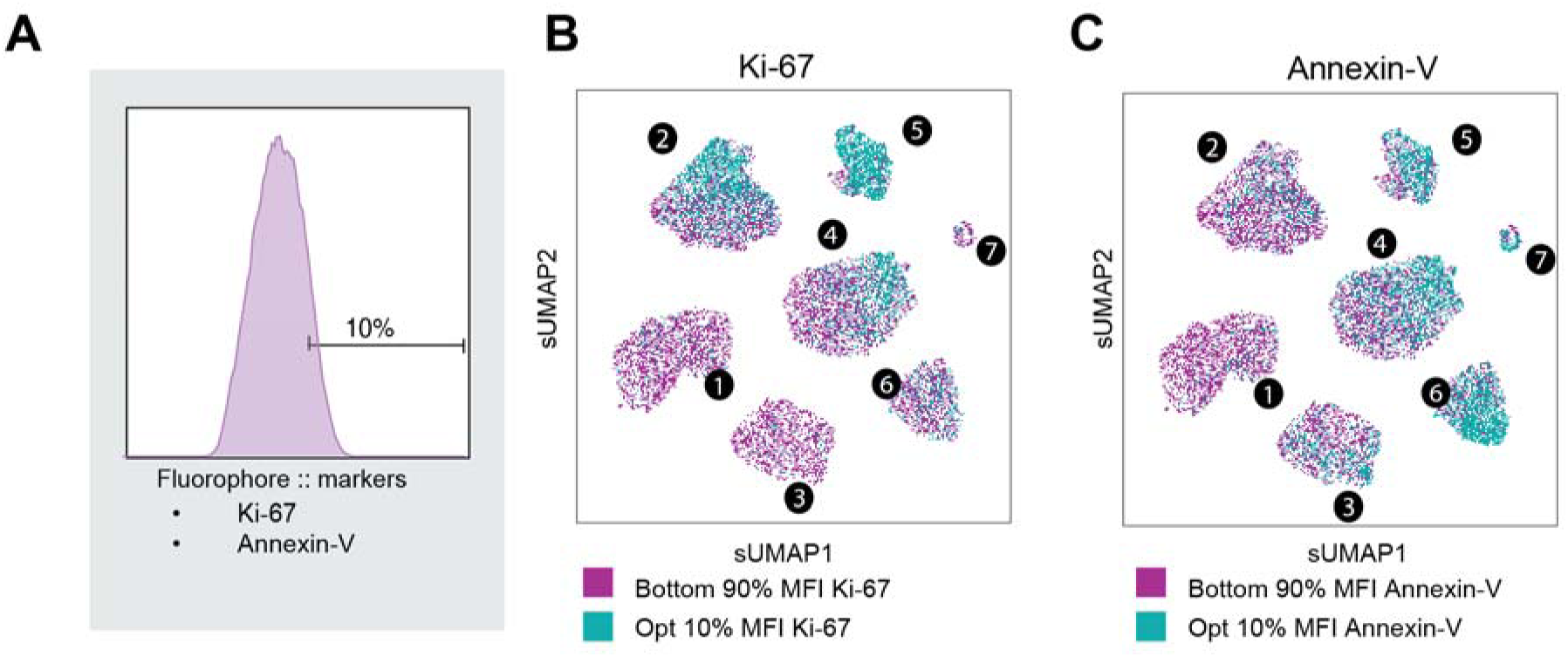
Fibroblast Subpopulations Exhibit Distinct Proliferative and Apoptotic Profiles. (A) Gating strategy for intracellular flow cytometry to quantify Ki-67 (proliferation marker) and Annexin-V (apoptosis marker) in fibroblasts, highlighting the top 10% MFI (mean fluorescence intensity) population used to define high-expressing cells. (B-C) UMAP visualization of fibroblast subpopulations, showing the distribution of cells with high (B) Ki-67 and (C) Annexin-V+ cells expression (teal) and remaining 90% (magenta) across seven clusters.

**Supplementary Table 1.**
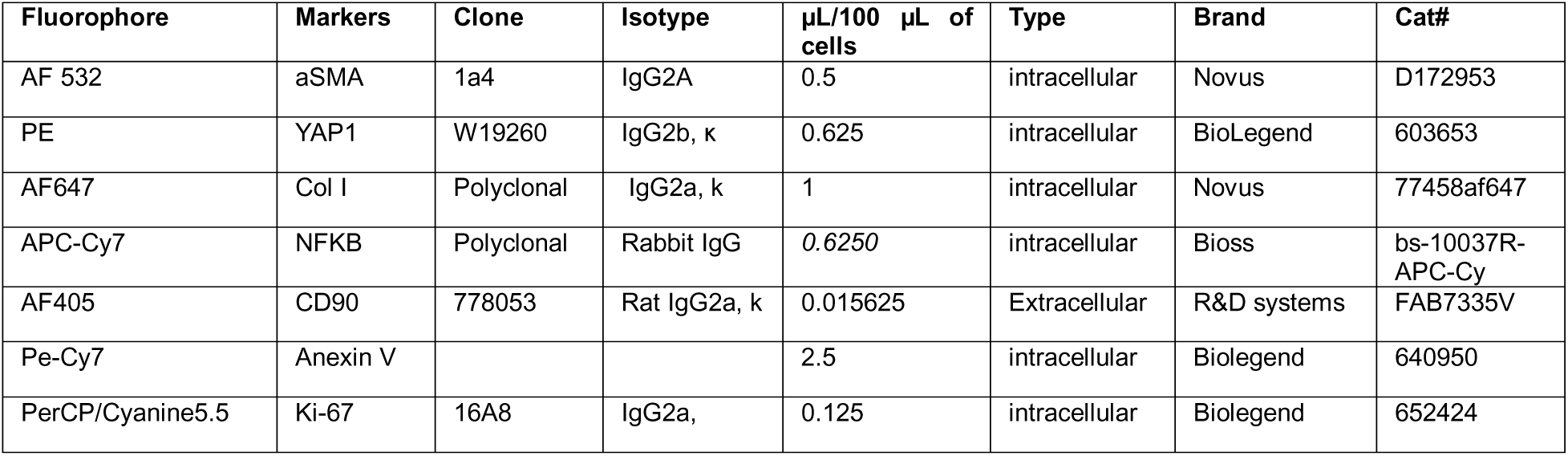
Flow cytometry markers.

**Supplementary Table 2.** Immunofluorescence markers.

| Fluorophore | Markers | Clone | Isotype | Brand | Concentration |
| --- | --- | --- | --- | --- | --- |
| FITC | aSMA | 1A4 | IgG2a | eBiosciences | (1:250) |
| AF647 | Col I | Polyclonal | IgG2a, $\kappa$ | Novus | (1:100) |
| APC-Cy7 | NFKB | Polyclonal | Rabbit IgG | Bioss | (1:100) |
| AF488 | YAP | Polyclonal | Rabbit IgG | Cell signaling | (1:200) |
| Rhodamine | Phalloidin |  |  | Abcam | (1:1000) |

